# A comparison of behavior paradigms assessing spatial memory in tree shrew

**DOI:** 10.1101/2023.05.16.540961

**Authors:** Cheng-Ji Li, Yi-Qing Hui, Rong Zhang, Hai-Yang Zhou, Xing Cai, Li Lu

## Abstract

Spatial cognition enables animals to navigate the environment. Impairments in spatial navigation are often preclinical signs of Alzheimer’s disease (AD) in human. Therefore, evaluating spatial memory deficits is valuable when assessing incipient AD in animal models. The Chinese tree shrew, a close relative of primates, possesses many features that make it suitable for AD research. However, there is a scarcity of reliable behavior paradigms to monitor changes in spatial cognition in this species. To address this, we established reward-based paradigms in the radial-arm maze and the cheeseboard maze for tree shrew, and tested spatial memory of a group of twelve male animals in both tasks, along with a control water maze test, before and after bilateral lesions to the hippocampus, the brain region essential for spatial navigation. Tree shrews memorized target positions during training, and their task performance improved gradually until reaching a plateau in all three mazes. After the lesion, spatial learning was compromised in both newly-developed tasks, whereas memory retrieval was impaired in the water maze. Furthermore, individual task performance in both dry-land paradigms depended heavily on the size of remaining hippocampal tissue. Notably, all lesioned animals displayed spatial memory deficits in the cheeseboard task, but not in the other two paradigms. Our results suggest that the cheeseboard task currently represents the most sensitive paradigm for assessing spatial memory in tree shrew, with the potential to monitor progressive cognitive declines in aged or genetically modified animals developing AD-like symptoms.

**Significance Statement:** Cognitive tests that monitor impairments in spatial memory play a crucial role in evaluating animal models with early-stage Alzheimer’s disease (AD). The Chinese tree shrew possesses many features suitable for an AD model, yet behavior tests assessing spatial cognition in this species are lacking. Here we developed novel behavior paradigms tailored to measure spatial memory in tree shrews and evaluated their sensitivity to changes in spatial learning by examining a group of hippocampus-lesioned animals. Our results indicate that the cheeseboard task effectively detects impairments in spatial memory and holds potential for monitoring the progressive cognitive decline in aged or genetically modified tree shrews that develop AD-like symptoms. This research may facilitate the use of tree shrew model in AD research.

## 1. Introduction

The ability to navigate through an environment is crucial for the daily life of animals. Early studies have demonstrated that lesions to the entorhinal-hippocampal circuit compromise an animal’s spatial learning ability (Squire, 1992; Steffenach et al., 2005). In human, damage to the entorhinal-hippocampal circuit is primarily observed in patients suffering from Alzheimer’s disease (AD), an age-related progressive neurodegenerative disorder (Braak and Braak, 1991; Adams et al., 2022). Although loss of episodic memory is a standard clinical diagnostic measure for AD (Dubois et al., 2014), deficits in spatial navigation are more sensitive in identifying at-risk individuals (Coughlan et al., 2018). Emerging data obtained from rodent models also reveal early AD pathophysiology in the entorhinal-hippocampal circuit, suggesting that navigation deficit can be a more sensitive cognitive marker for this disease (Fu et al., 2017; Jun et al., 2020; Ying et al., 2022). In summary, spatial navigation abilities are the most vulnerable to damage in the entorhinal-hippocampal circuit, and can serve as a cognitive marker for incipient AD, both in human patients and in animal models.

Chinese tree shrew (*Tupaia belangeri*) is a squirrel-like mammal currently placed in the order Scandentia (Zheng et al., 2014). Due to its short reproductive cycle and phylogenetically close affinity to primates (Xu et al., 2012; Fan et al., 2013; Fan et al., 2019; Ye et al., 2021), tree shrew has been used as experimental models alternative to primates, in a variety of human diseases, including infections, cancer (Cao et al., 2003; Xiao et al., 2017; Yao, 2017; Li et al., 2018), and neuropsychiatric disorders (Fuchs, 2005; Zambello et al., 2010; Pryce and Fuchs, 2017; Ni et al., 2020; Savier et al., 2021). The amino acid sequence of beta-amyloid (Aβ) in tree shrew is identical to that in human (Pawlik et al., 1999), and the expression patterns of AD-related genes are similar (Fan et al., 2018; Ye et al., 2021). Additionally, Aβ deposition and tau over-phosphorylation, two key molecular markers for AD, have been observed in the brains of aged tree shrews (Yamashita et al., 2010; Yamashita et al., 2012; Fan et al., 2018), suggesting that AD may occur naturally in those animals, similar to non-human primates (Paspalas et al., 2018; Li et al., 2022). Moreover, studies have shown that intracerebroventricular injections of Aβ fragments into the tree shrew brain result in profound molecular, cellular, and cognitive changes (Lin et al., 2016; Wang et al., 2020), which closely resemble human AD pathology (Masters et al., 2015). Collectively, these pioneer studies suggest that tree shrew may serve as a promising animal model for future AD research.

Cognitive impairment is a critical indicator for a valid AD model (Chen and Zhang, 2022). A number of behavioral paradigms have been developed to study cognitive functions in tree shrew, such as associative learning (Ohl et al., 1998), object recognition (Khani and Rainer, 2012; Nair et al., 2014), contextual fear conditioning (Shang et al., 2015) and visual discrimination (Mustafar et al., 2018). However, these tasks are not specifically designed to evaluate tree shrew’s memory of location in space, an important measure for spatial cognition. Although tree shrews have been tested in the water maze for loss of spatial memory (Wang et al., 2020), this paradigm is only sensitive for animals with severe hippocampal damages (Moser et al., 1993; Moser et al., 1995). Taken together, these behavioral tests are not suitable for detecting mild spatial cognitive impairment representing early stage of AD.

The aim of this study was to establish robust behavioral paradigms that can detect small changes in spatial memory in tree shrew, so that the progressive declines in spatial cognition can be reliably monitored in animals developing AD. Here we developed tasks in the radial-arm maze and the cheeseboard maze for tree shrew, and compared their sensitivity to spatial memory loss in hippocampus-lesioned animals with the pre-established water maze test.

## 2. Materials and methods

### 2.1. Subjects

The experiments were conducted at the Kunming Institute of Zoology (Kunming, China), in accordance with the guide for the care and use of laboratory animals, approved by the Institutional Animal Care and Use Committee of Kunming Institute of Zoology (KIZ-IACUC-TE-2022-02-001). Fifteen adult male Chinese tree shrews (*Tupaia belangeri*), aged 13-15 months and weighing 110-150 g at the start of the experiment, were initially included in the experiments (TS094-TS108). These animals had no prior experience in any behavioral test and were housed individually in wire cages (44×38×35 cm), each equipped with a dark nest box (36×16×20 cm), under standard laboratory conditions (temperature: 20-26, relative humidity: 40-60%) on a 12 h light/dark schedule. Testing was conducted in the light phase. Three animals were removed before test start due to health issues (TS098 & TS107, vulnerable to food restriction) or aversion to tests (TS105, untrainable). The remaining twelve animals were tested in the radial-arm maze, the cheeseboard maze, and the water maze sequentially, with one-week interval between two tasks (**Fig. 1 A-F**). Tree shrews were mildly food restricted during test period in the radial-am and cheeseboard mazes (kept at 85–90% of free-feeding body weight), otherwise they were fed *at libitum* (**Fig. 1 G**). Twelve tree shrews received neurotoxic lesions to the hippocampus after the first round of behavior tests, and eleven of them were later tested sequentially in the same tasks. One animal (TS104) died during surgery, possibly due to isoflurane overdose and was excluded from pre-and post-lesion comparison analysis. The animals’ running speed was slightly higher in the cheeseboard maze after bilateral lesions to the hippocampus (**Fig. 1 H**), while their swimming speed in the water maze was not altered (**Fig. 1 I**).

**Figure 1.**
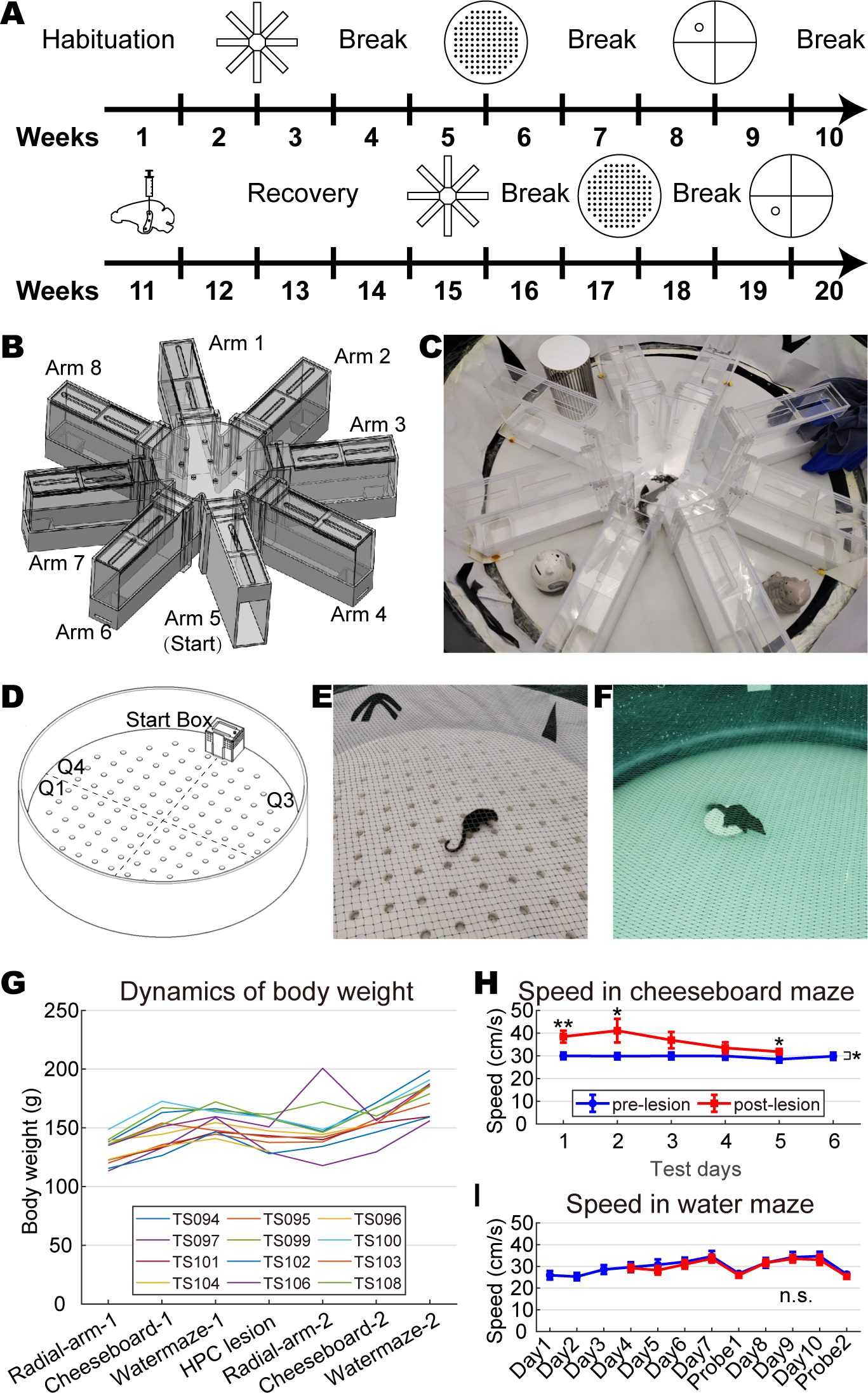
Experimental design and behavior apparatus. **A**, Timeline of the experiments. **B**, Design of the tree shrew radial-arm maze, with transparent walls and lids to facilitate the animal’s perception of environmental cues. The start arm (arm 5) contains a large opening that connects to a transfer box. **C**, The radial-arm maze is surrounded by various visual cues. The tree shrew in the maze center is performing the task. **D**, Schematic of the cheeseboard maze. **E**, The cheeseboard maze is decorated with visual cues. **F**, A tree shrew seated on the platform in the water maze (visible platform during pre-training). **G**, The body weight of tree shrews at the time each task/procedure began. Colored lines represent data from individual animals. Their bodyweight remained largely constant with two exceptions: TS097 and TS106 had a dramatic decrease and increase, respectively after the lesion. **H**, Tree shrews’ running speed (mean ± SEM, filtered with 2.5 cm/s threshold) in the cheeseboard maze before and after the hippocampal lesion. They ran faster after lesion, especially in the first two days (two-way ANOVA for repeated measures, 11 animals, test day: F (4) = 4.053, p = 0.008, η^2^ = 0.288; lesion: F (1,4) = 7.833, p = 0.019, η^2^ = 0.439; lesion × test day: F (1,4) = 2.699, p = 0.044, η^2^ = 0.213). **I**, Tree shrews’ swimming speed (mean ± SEM, filtered with 2.5 cm/s threshold) in the water maze did not differ before and after the lesion (two-way ANOVA for repeated measures, test day: F (6) = 7.138, p < 0.001, η^2^ = 0.417; lesion: F (1) = 1.025, p = 0.335, η^2^ = 0.093; lesion × test day: F (1,6) = 0.269, p = 0.949, η^2^ = 0.026). n.s., not significant, *, p < 0.05, **, p < 0.01.

### 2.2. Experimental Design

The experimental design of this study is shown in **Fig. 1 A**. In brief, a group of twelve male adult tree shrews were first trained and tested sequentially in the radial-arm maze, the cheeseboard maze and the water maze, which are behavior paradigms specifically designed to assess their spatial memory. In order to obtain their learning profiles in each behavior test, the animals were tested consecutively until their learning curves plateaued. All animals were next subjected to neurotoxic procedures to induce bilateral lesions to the hippocampus to impair their abilities in spatial memory. After recovery, the same tree shrews were finally re-tested in these cognitive tasks to measure their loss of spatial memory. This experimental designed not only allows each animal to serve as its own control so that interindividual variability can be avoided, but also enables us to compare these behavior paradigms in the same animal. ANOVA for repeated measures and tests for paired samples were used to compare spatial learning before and after the lesion. Details of the above experiments are described in following sections.

### 2.3. Behavior apparatus

#### 2.3.1. Radial-arm maze

The radial-arm maze was originally designed for assessing working memory and reference (spatial) memory in rats (Olton and Samuelson, 1976). The maze, which was 128 cm in diameter and 28 cm in height, was constructed by SANS Biological Technology using white plastic (floor, doors, lower 1/3 of walls) and transparent plexiglass (upper 2/3 of walls and lids) (**Fig. 1 B**). It was positioned in a well-lit room (3.0×2.5 m) and placed on a platform 36 cm above the ground, surrounded by numerous visual cues. The maze comprised a center region (28 cm in diameter) and equally spaced 8 arms (50 cm in length and 10 cm in width). A small cake reward, placed in a square bowl (9.0×9.0×5.2 cm) positioned at the end of each arm, was not visible to the animal from the other end of the arm. Order cues were obscured by large pieces of cake placed near the end of each arm outside the maze. Prior to the beginning of testing, the animals were pretrained to enter all eight arms to receive cake rewards. During testing, the eight arms were separated into one closed start arm (arm 5), three baited open arms (arms 2, 4, 7 before lesion; arms 3, 6, 8 after lesion) and four un-baited open arms. At the start of each test day, each animal was transferred into the start arm, and allowed to acclimate for a few minutes. The start arm was covered by a towel when rewards were placed, so that the animal waiting inside had no hint of the reward locations. For each trial, tree shrew walked into the center region after the start arm was opened (start signal), and then entered the remaining seven arms to find rewards (**Fig. 1 C**). Trial ended when all three rewards had been consumed or after two minutes. Once the start arm was reopened (stop signal), the animal returned to the start arm (manually guided if necessary) to receive a fourth cake reward and then waited for one minute for the next trial began. Testing for each animal concluded at 25 trials or 50 minutes, whichever occurred first. Urine marks were removed with bleach after each trial, and the entire maze was cleaned thoroughly with 70% alcohol once the testing for each animal was completed.

#### 2.3.2. Cheeseboard maze

The cheeseboard maze was originally designed for navigation studies in rat (Dupret et al., 2010). The maze was constructed with steel walls and a white plastic floor, measuring 150 cm in diameter and 50 cm in height. The maze was placed on a platform 36 cm above the floor, in a well-lit room (3.0×2.5 m), decorated with numerous visual cues to help the animals orient themselves. The maze floor was divided into four quadrants, each containing 30 equally spaced wells (3.0 cm in diameter, 3.5 cm in depth, spaced in 10.0 cm). A transparent start box (22×16×20 cm) was positioned at the boundary of quadrants 3 and 4 near the wall (**Fig. 1 D**). Prior to the everyday test, the animals were pretrained to navigate in the maze and consume cake rewards placed in three distant wells. During the everyday test, three cake rewards were placed in wells that met the three criteria: a) they were not chosen on the previous test day; b) the distance between any two wells was greater than 30 cm, and c) the area of the triangle formed by the three reward wells was 11-12 dm^2^. Locations of reward wells before lesion were: test day 1: [−3, 4; −1, −3; 3, −1]; test day 2: [2, 5; −3, 5; −1, −2]; test day 3: [4, 1; −2, 5; −4, 2]; test day 4: [3, 1; −4, 3; −4, −2]; test day 5: [3, −1; −1, 4; −4, −1]; test day 6: [3, 4; −4, 3; −2, −2]. Locations of reward wells after hippocampal lesion were: test day 1: [3, 4; 1, −3; −3, −1]; test day 2: [−2, 5; 3, 5; 1, −2]; test day 3: [−4, 1; 2, 5; 4, 2]; test day 4: [−3, 1; 4, 3; 4, −2]; test day 5: [−3, −1; 1, 4; 4, −1]. The spatial configuration of reward wells before and after the lesion was symmetrical with respect to the vertical midline. To prevent tree shrew from relying on local cues to solve the task, rewards were not visible to the animal, and order cues were obscured using a layer of cake crumbs that were evenly spread beneath the floor. At the start of each test day, each animal was transferred into the start box, and allowed to acclimate for a few minutes. The start box was covered by a towel when rewards were placed, so that the animal waiting inside had no hint of the reward locations. During each trial, the tree shrew was free to navigate the maze to locate and consume three cake rewards after the start box was opened (start signal) (**Fig. 1 E**). Trial ended when all three rewards had been consumed or after two minutes. Once the start box was reopened (stop signal), the animal returned to the start box (manually guided if necessary) to receive a fourth cake reward and then waited for one minute for the next trial began. Testing for each animal concluded at 25 trials or 50 minutes, whichever occurred first. Urine marks were removed with bleach after each trial, and the entire maze was cleaned thoroughly with 70% alcohol once the testing for each animal was completed.

#### 2.3.3. Water maze

The water maze was originally designed to access spatial memory in rats (Morris, 1984). The steel swimming pool was 150 cm in diameter and 50 cm in depth, with a featureless, black inner surface (SANS Biological Technology). The maze was positioned in a well-lit room (3.0×2.5 m) with numerous visual cues. The pool was filled to a depth of 20 cm with water maintained at a temperature of 18°C, to which 60 g of titanium dioxide powder was added. A white metal platform measuring 14 cm in diameter was submerged 1.5 cm below the water surface in the center of quadrant 2 (before lesion) or quadrant 3 (after lesion) during training. During pretraining (day 0), the water level was lowered 3 cm to make the platform visible to the animal (**Fig. 1 F**). Subsequently (days 1-7), the tree shrews were trained for four trails per day, spaced 30 minutes apart. Each trial was begun by releasing the animal into the water with its face oriented towards the pool wall, at one out of four quadrants selected at random. If the tree shrew failed to locate the platform within 60 seconds, it was manually guided onto it. The animal was always allowed to stay on the platform for 20 seconds. During the spatial probe test, the tree shrews were released from quadrant 4 (before lesion) or quadrant 1 (after lesion), and their swimming was tracked for a duration of 60 seconds. Upon completion of each trial or test, the animals were dried with a towel and allowed to rest in their nest boxes beside a heater.

### 2.4. Behavioral data collection

The animals’ behaviors during training and testing were monitored by an overhead camera mounted 1.9 m above the mazes. Videos were captured with a resolution of 640×480 px, at a sampling rate of 50 Hz, in .avi format, using Captura software (Version 9.0.0, open-source). The movements of the tree shrews were then tracked off-line using DeepLabCut software (RRID: SCR_021391), which is an open-source tool that uses transfer learning with deep neural networks for markerless pose estimation (Mathis et al., 2018). To train the network model for a given animal, 600-1000 frames from 4-7 videos were manually labelled to identify the snout, eyes, and tail base when necessary, and trained 900,000 times to ensure accuracy. The trained model was then used to automatically label all videos for the animal. Any labelling errors identified in DeepLabCut outputs (time-position series of labelled points), such as discontinuities and sudden jumps, were manually corrected using custom MATLAB scripts.

### 2.5. Behavior analyses

Tree shrews’ performance in the behavior tests was assessed using custom MATLAB (RRID: SCR_001622) scripts, which were designed to analyze data obtained from the DeepLabCut. To ensure the accuracy of the results, the output data from the MATLAB scripts was verified by manual inspection of the original trial videos.

#### 2.5.1. Radial-arm maze

Tree shrew’s snout positions were used to analyze their task performance. A valid arm visit was defined as snout position greater than 35 cm into the arm.

Working memory error rate for each trial was determined as:

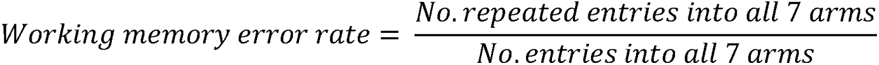

Reference (spatial) memory error rate in each trial was determined as:

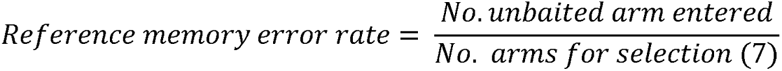

In instances where the animal was unable to locate all three rewards before the end of the trial, only arm entries within the first two-minute time frame were included for analysis. A trial was deemed correct if all three rewards were found without erroneous arm visits.

#### 2.5.2. Cheeseboard maze

Both tree shrew’s snout and head (midpoint of left and right eyes) positions were used as tracked variables. A valid visit to a reward well was defined as a snout position in the well longer than 0.1 seconds. A trial was deemed successful if all three rewards were found within two minutes. Only trials that occurred after the first successful trial on each test day were included in subsequent analyses.

Routescores were used to evaluate tree shrews’ spatial memory of reward locations:

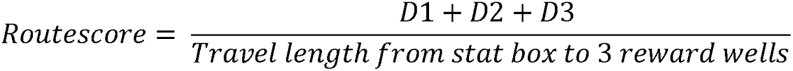

Where D1 is the distance between the start box and the first reward well visited, D2 is the distance between the first and second reward wells, and D3 is the distance between the second and third reward wells. If a trial was unsuccessful, the routescore was defined as 0. Test days on which the animal performed five or fewer successful trials were considered unmotivated and were excluded from the analyses.

#### 2.5.3. Water maze

Tree shrew’s head positions were used to assess their performance in the water maze. Escape latency and distance were calculated as the time and travel length, respectively, that the animal took before climbing onto the platform. A successful trial was defined as the animal found the platform. The escape latency was defined as 60 seconds in unsuccessful trials, and the escape distance was defined as the travel length during this time. In the spatial probe test, a valid crossing of the platform was defined as the animal spending more than 0.1 s in the platform region. The occupancy of the target quadrant was evaluated by the proportion of time or distance in the quadrant where the platform used to locate, which was calculated as:

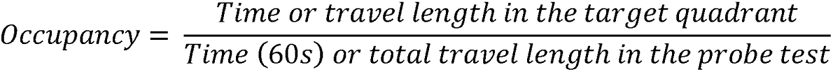

### 2.6. Surgery

Twelve tree shrews were subjected to a neurotoxic procedure to induce lesions in their hippocampi. The animals were anesthetized with isoflurane (airflow: 0.8-1.0 L/min, 0.5-3% isoflurane mixed with oxygen, adjusted according to physiological monitoring, RWD Life Science). Their body temperature was maintained at approximately 38 using a heating pad underneath the body with a closed-loop controller connected to a rectal temperature probe. Upon induction of anesthesia, Metacam (2 mg/ml, 1mg/kg) and Baytril (50 mg/ml, 5 mg/kg) were injected subcutaneously, the animal’s head was then fixed in a stereotaxic frame (RWD Life Science) with custom-designed gas mask. A local anesthetic (2% Lidocaine, 200 μl) was applied under the skin before making the incision. Both temporalis muscles were gently detached from the skull and slightly moved such that infusion holes could be drilled in the skull above the hippocampus. The neurotoxin (colchicine, MedChemExpress, CAS: 64-86-8, 0.6 mg/ml) was dissolved in sterile phosphate-buffered saline (pH7.2) and injected using a sharp 5-μl syringe (Hamilton) mounted to the stereotaxic frame (with the opening of the needle directed caudally to the animal). Volumes of 1.0 μl of colchicine solution were infused at a speed of 10 μl /h using a micro pump (KD Scientific), at three stereotaxic positions in the left and right hippocampi, using bregma as a reference for infusion coordinates. The infusion coordinates were: ① dorsal: AP = +4.5 mm, ML = ±5.0 mm, DV = −4.0 mm, ② intermediate: AP = +4.5 mm, ML = ±7.0 mm, DV = −7.0 mm, and ③ ventral: AP = +2.5 mm, ML = ±4.5 mm, DV = −11.2 mm. At each position, the needle was first lowered to the cranium below the injection site and then retracted for 0.1 mm. The injection started 1 minute after the needle was positioned. After the injection, the needle was left in place for 10 minutes to allow absorption. When the injections were completed, the skull was cleaned with saline solution and the skin was sutured. The animals received post-operative care, including soft food, analgesics, and antibiotics for three days, followed by a three-week recovery period with free food and water, before the second round of behavior tests were performed.

### 2.7. Histology

At the end of the experiment, the tree shrews were deeply anesthetized with overdose isoflurane, and perfused transcardially with 0.9% saline, followed by 10% formalin. After extraction from the skull, the brains were post-fixed in 10% formalin overnight and subsequently in a sequence of 20%, 30%, and 30% sucrose for 2-3 weeks until sectioning. Coronal sections (40 µm) were cut on a cryostat (KEDEE, KD-2950) and stained with cresyl violet. Cresyl violet staining was carried out on sections mounted on microscope slides (CITOGLAS). The sections were first dehydrated in graded ethanol baths (70%, 80%, 90%, 100%, 100%, and 100%), cleared in turpentine oil and then rehydrated in a reverse direction in the same set of ethanol baths before they were stained in a 0.1% cresyl violet solution (Sigma Aldrich, CAS: 10510-54-0, Cat# C5042-10g) for 5-10 minutes. The staining was differentiated by dipping the sections in a solution consisting of 70% ethanol and 0.5% acetic acid, and after this the sections were dehydrated again in ethanol baths and cleared in turpentine oil before they were cover-slipped with neutral balsam. Finally, the Nissl-stained brain sections were imaged under a brightfield microscope, with a 10×, 0.4 NA objective (Olympus BX61), using the accompanying software OlyVIA V3.2 (Build 21633, RRID: SCR_016167).

### 2.8. Lesion quantification

To quantify the extent of hippocampal damage in each lesioned tree shrew, cell loss in each equally spaced brain section (120 μm) were identified. Regions of healthy hippocampal tissue (including dentate gyrus, CA3, CA2, CA1 and subiculum) in each coronal section were manually labelled and measured by the ImageJ software (RRID: SCR_003070) (Schneider et al., 2012). The volume of healthy hippocampal formation was calculated by summing all labelled regions and multiplying by the space between sections. The percentage of hippocampal damage was expressed as the amount of lesioned tissue (volume of a standard hippocampus subtracted by healthy hippocampal volume) in proportion to the volume of the standard hippocampus (78.10 mm^3^). This approach allowed us to quantitatively assess the severity of hippocampal damage in each tree shrew and compare the results across individuals.

### 2.9. Statistical Analysis

The results are presented as mean ± standard error of the mean (SEM). One-way ANOVA for repeated measures was applied to compare the learning curves over days before or after the hippocampal lesion. Task performance before and after the lesion was assessed with two-way ANOVA for repeated measures, using a 2 × N design with both lesion and test day as within-subject factors. Bonferroni corrections were applied for multiple comparisons to reduce the likelihood of false positives. Wilcoxon signed-rank test and paired t-test were used to compare two groups of related measures. One sample t-test was used to compare the mean of measures with an expected value. Pearson correlation was used to evaluate the relationship between task performance and damage in the hippocampus. Significance level was set at p < 0.05, and was given for two-tailed tests. All statistical analyses were performed using SPSS statistics software (IBM, RRID: SCR_002865) version 25.

## 3. Results

### 3.1. The radial-arm task for tree shrew

Tree shrew is a skittish and agile animal, which poses challenges for behavior experiments. To overcome these challenges, we first habituated the naïve animals to the lab environment and experimenters for one week and then trained and tested them in the novel radial-arm maze that was specifically tailored to the behavioral characteristics of tree shrews (**Fig. 1 A-C**). In contrast to the rodent version, our radial-arm task necessitated that the tree shrews entered the center region from a designated start arm (arm 5) after trial started, and returned to the same arm after completing each trial. The design of the maze allowed us to guide the animals to their destinations without provoking agitation. Cake rewards were placed in arms 2, 4, and 7, while the remaining four arms (1, 3, 6, and 8) were not baited (**Fig. 1 B, Fig. 2 A**).

**Figure 2.**
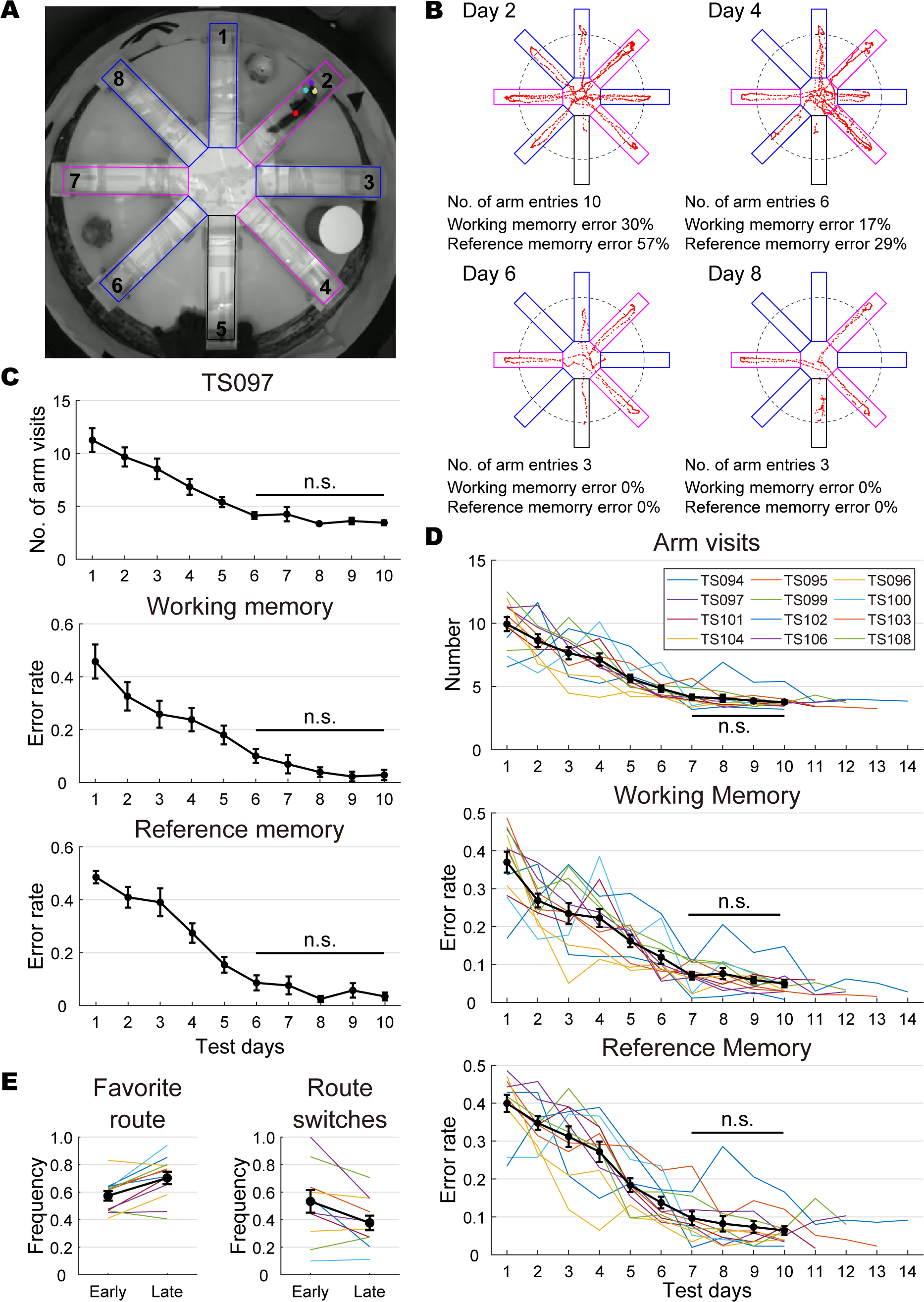
Performance of tree shrews in the radial-arm maze prior to hippocampal lesion. **A**. Camera footage of a tree shrew performing the radial-arm task. The colored boxes denote the arms of the maze for subsequent analysis. Black: the start arm; magenta: baited arms; blue: un-baited arms. Arm IDs are indicated by numbers. The colored dots on the animal are automatically labelled marks from DeepLabCut. **B**, Representative trials performed by TS097 in the radial-arm task. The red dots indicate the location of tree shrew’s snout. The dashed black circle denotes the threshold of a valid arm visit, which is counted when the snout position goes beyond the threshold. The colored boxes are the same as in panel **A**. **C**, TS097’s task performance in the radial-arm maze during the first ten test days, assessed by the number of arm visits (mean ± SEM) upon trial completion (top), error rates in working memory (middle), and error rates in reference memory (bottom) (one-way ANOVA for repeated measures, arm entry: F(9) = 43.406, p < 0.001, η^2^ = 0.644; working memory: F(9) = 28.115, p < 0.001, η^2^ = 0.539; reference memory: F(9) = 54.623, p < 0.001, η^2^ = 0.695). **D**, Tree shrews’ performance in the radial-arm maze plateaued after day 7, as shown by the number of arm visits (top), error rates in working memory (middle) and error rates in reference memory (bottom). The black traces represent the mean ± SEM, while the colored lines indicate data from individual animals. **E**, Tree shrews’ preference for certain sequences after intensive training, measured by the frequency of favorite route (left) and frequency of route switches (right). The color code is the same as in **D**. n.s., not significant.

During repeated training in the radial-arm maze, the tree shrews demonstrated a decrease in the number of visits to un-baited arms and a reduction in the number of re-visits to baited arms over time (**Fig. 2 B**). To measure the frequency of erroneous arm visits, we defined an arm visit as the nose position passing > 35 cm into the arm, which corresponds to the commonly used 0.3× arm threshold for rodents (four paws passing > 15 cm, plus a body length of 20 cm) (Masuda et al., 1994). With continued training, the tree shrews showed a progressive decrease in the number of arms visited before consuming three rewards, as well as a decline in both working memory error rates (frequency of repeated visits to arms) and reference memory error rates (frequency of visits to un-baited arms, which is spatial memory dependent) (**Fig. 2 C**). To determine the learning profile of tree shrews in the radial-arm paradigm, each animal was consecutively tested for at least ten days, until both working memory and reference memory error rates were below 0.15, which is a common criterion used in rodent behavior experiments (Igarashi et al., 2014). After meeting this criterion, the animals were further tested for three more days. Although there were noticeable individual differences in task acquisition, once the criterion was met, tree shrews’ task performance remained stable in the following days, with error rates fluctuating slightly but never exceeding 0.15. Thus, we defined that the tree shrews had learnt the radial-arm task once both error rates were lower than 0.15. On average, tree shrews learned the task on the 6.83 ± 0.56 day (mean ± SEM).

We next quantified the performance of twelve tree shrews in the radial-arm maze over the period of ten days, with three measures: the number of arm visits upon trial completion, as well as the error rates for working memory and reference memory. We observed a consistent decrease in all three measures over test days, which was statistically significant (one-way ANOVA for repeated measures, 12 animals, arm visits: F(9) = 39.904, p < 0.001, η^2^ = 0.784; working memory: F(9) = 39.476, p < 0.001, η^2^ = 0.782; reference memory: F(9) = 51.746, p < 0.001, η^2^ = 0.825). The performance plateaued after 6 days of training, which we confirmed through post-hoc analysis with Bonferroni correction (all p values ≥ 0.795) (**Fig. 2 D**). We also noticed that tree shrews tended to develop a preference for a specific route when visiting the baited arms (stereotyped route pattern in consecutive trials), especially after they had mastered the task. We quantified this stereotypy by measuring the frequency of the favorite route in correct trials (trials without any erroneous arm visits) and the frequency of route switches (the proportion of correct trials in which a different route from the previous trial was selected). We found that the route preference increased significantly from 0.573 ± 0.035 in the early stage (test days 1-7), to 0.703 ± 0.045 in the late stage (test days 8-10) (paired t-test, t(11) = 3.948, p = 0.002), while the frequency of switches decreased (from 0.533±0.082, to 0.376 ± 0.053, t(11) = 2.747, p = 0.019) (**Fig. 2 E**), raising the possibility that the tree shrews may have developed alternative strategies to complete the task after extensive training. Based on our findings, we concluded that a seven-day test period was optimal for the tree shrew radial-arm task.

### 3.2. The cheeseboard task for tree shrew

Tree shrews were next trained and tested in the cheeseboard maze (**Fig. 1 A**). Unlike the radial-arm task, where rewards were always baited in the same arms, a different set of reward wells was intentionally selected for the cheeseboard task each day (see methods), which required daily memory updates of goal locations. The distances between the three baited wells were kept comparable to maintain consistent difficulty levels across days. With previous experience in the radial-arm maze, the tree shrews were more collaborative in the cheeseboard maze. They were easily lured back to the start box with a cake reward at trial end. The animals were tested in the cheeseboard maze for six consecutive days after learning to find rewards from three distant wells (**Fig. 3 A**).

**Figure 3.**
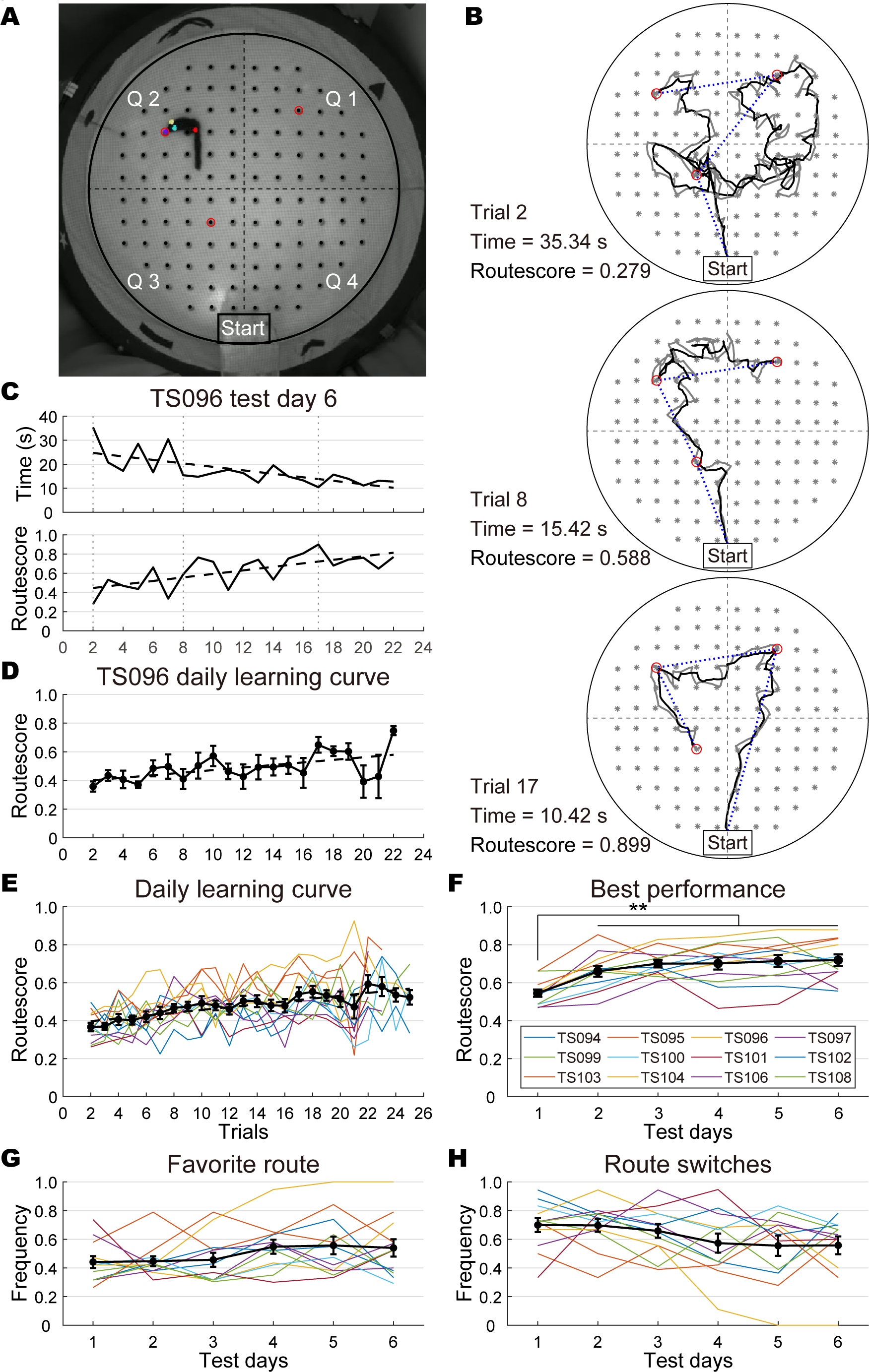
Performance of tree shrews in the cheeseboard maze before the lesion. **A**. Camera footage of a tree shrew performing the cheeseboard task. The maze is partitioned into four quadrants using dashed lines. Baited wells were indicated by red circles. The colored dots on the animal are automatically labelled marks from DeepLabCut. **B**, Representative trials performed by TS096 in the cheeseboard maze on test day 6. The black and grey traces show the trajectory of tree shrew’s head and snout, respectively. The positions of all 120 wells are marked with grey stars. The three baited wells are marked with red circles. The blue dotted lines show the distance between the start box and the three baited wells, which was used to calculate the routhscore. **C**, TS096’s learning curve in the cheeseboard maze on test day 6, evaluated by time spent (top) before consuming three rewards and routhscore (bottom). The dashed lines indicate the learning trend. The vertical dotted lines refer to trials shown in panel **B**. **D**, TS096’s daily learning curve (mean ± SEM) in the cheeseboard maze, averaged across six test days (one-way ANOVA for repeated measures, F(20) = 2.566, p = 0.001, η^2^ = 0.339). **E-F**, The daily learning curve (**E**) and best performance (**F**) of individual tree shrews in the cheeseboard maze, averaged across six test days and five best trials on each test day, respectively. Data from individual animals are shown in color lines, while the mean ± SEM of data from twelve animals is represented by the black trace. **G-H**, Tree shrews’ route preference over test days, measured by frequency of favorite route (**G**) and frequency of route switches (**H**). Only TS104 displayed a clear preference over time. The same color code is used as in panel **F**. **, p < 0.01.

During the six-day test, the time each tree shrew spent, and the distance travelled to find the rewards, were reduced in later trials (**Fig. 3 B**). Since the animals could visit the baited wells in different sequences and their running speed varied from trial to trial, a new measure named “routescore”, was used to evaluate tree shrew’s memory of target locations in the maze. Routescore was defined as the ratio between ideal route to the three rewards and travel distance in real, with a higher routescore indicating a more accurate spatial memory. On each test day, routescore increased gradually with more trials, accompanied by a parallel decrease in trial time (**Fig. 3 C**), indicating an improvement in their memory of reward locations over training. A similar trend of routescore increase was observed when the six test days were combined (**Fig. 3 D**). The analysis of daily learning curves from all twelve animals showed that their performance in the cheeseboard maze improved progressively within 25 trials (one-way ANOVA for repeated measures, 12 animals, F(23) = 5.608, p < 0.001, η^2^ = 0.338) (**Fig. 3 E**).

We noticed that in some animals, routescores decreased near the end of testing on each day, possibly due to diminished motivation to complete the task. Thus, the mean value of best five trials (those with highest routescores), instead of the last five trials, was calculated to represent the best performance an animal can reach on every day test. We found that although some animals showed slight fluctuations in performance over the test days, the group data indicated a rapid increase in performance that stabilized after the second day of training (one-way ANOVA for repeated measures, 12 animals, F(5) = 13.014, p < 0.001, η^2^ = 0.542, post-hoc analysis with Bonferroni correction, test day 1 was significantly lower than other days, all p values ≤ 0.003) (**Fig. 3 F**), while the mean running speed of the animals remained unchanged over days (one-way ANOVA for repeated measures, F(5) = 0.737, p = 0.599, η^2^ = 0.063, n = 12) (**Fig. 1 H**). We also investigated whether the animals developed any stereotypic behavior during the training sessions, as this could potentially affect their performance. We examined the frequency of the preferred route taken by each animal and the frequency of route switches, but found no significant changes over the test days for most animals (one-way ANOVA for repeated measures, 12 animals, preferred route: F(5) = 1.733, p = 0.142, η^2^ = 0.136; route switches: F(5) = 2.101, p = 0.079, η^2^ = 0.160) (**Fig. 3 G-H**). These results suggest that the contribution of running speed and route preference to task performance was minimal, and that tree shrews’ performance in the cheeseboard maze was stable after two days of training.

### 3.3. The water maze test for tree shrew

The tree shrews were trained and tested in the water maze after they had experienced the dry-land mazes (**Fig. 1 A**). Extensive training in previous tests allowed the animals to be manually taken out from their nest boxes before each trial, and carried from the platform and wiped dry with towels after each trial. Unlike in previous tasks, where each tree shrew had custom pretraining until they met the criterion for everyday tests (see methods), in the water maze test, all animals underwent the same pretraining, training and testing procedures, following the rodent protocols (Vorhees and Williams, 2006).

Twelve tree shrews were first pre-trained with a visible platform on day 0, followed by seven days of training with a hidden platform (days 1-7), and then their spatial memory was tested without the platform (probe 1). Different from rats, tree shrews swam vigorously to keep their heads above water in the swimming pool, implying their swimming abilities were not as good as rats’. It was difficult for them to keep swimming for longer than one minute. Therefore, a 60-second trial limit was applied for the water maze test instead of two-minute limits for the dry-land tasks. During the first stage of training, tree shrews gradually learned to search for the hidden platform (**Fig. 4 A**), and they managed to locate the hidden platform after a few days (**Fig. 4 B**). This was indicated by the decrease in time spent and distance travelled before climbing onto the platform (**Fig. 4 C**). The success rate on each training day, defined as the proportion of successful trials (where the animal found the hidden platform), increased monotonously from training day 1 to day 7 (**Fig. 4 D**), accompanied by a parallel increase in swimming speed (one-way ANOVA for repeated measures, n = 12, F(6) = 7.723, p < 0.001, η^2^ = 0.412) (**Fig. 1 I**). The spatial acquisition of the animals as a group during first stage of training in the swimming pool, reflected by escape latency and distance, which were defined respectively as time spent and distanced travelled before locating the platform, decreased monotonically (one-way ANOVA for repeated measures, n = 12, latency: F(6) = 16.671, p < 0.001, η^2^ = 0.602; distance: F(6) = 11.512, p < 0.001, η^2^ = 0.511) (**Fig. 4 E**).

**Figure 4.**
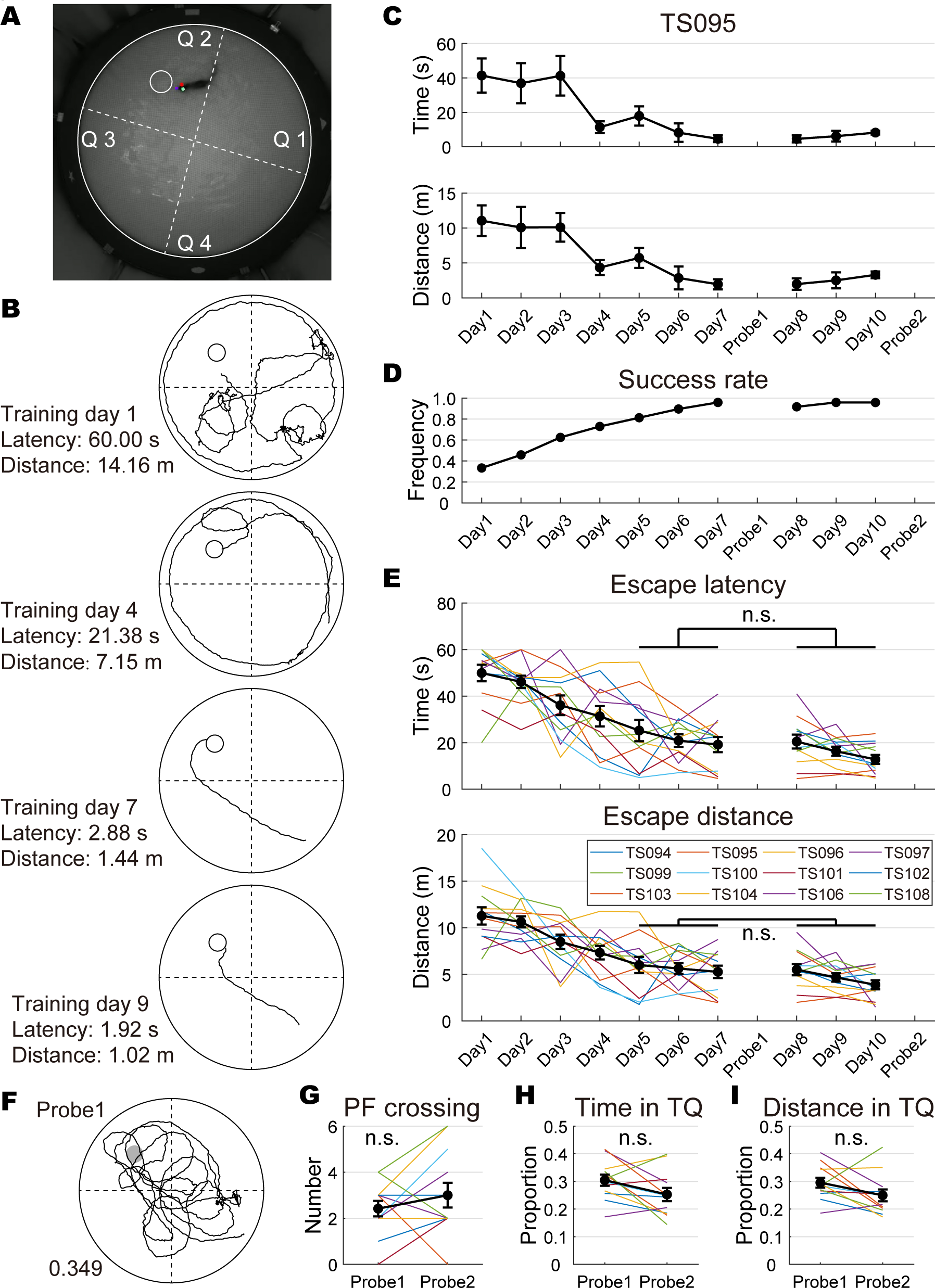
Performance of tree shrews in the water maze before lesion. **A**. Camera footage of a tree shrew performing the water maze test. The location of the platform is indicated by the smaller white circle and the maze is divided into four quadrants by dashed lines. The colored dots on the animal are automatically labelled marks from DeepLabCut. **B**, Representative training trials performed by TS095. The animal was released from the opposite quadrant. The trajectory of the animal’s head is indicated by black traces. The platform is represented by a smaller black circle. **C**, The performance of TS095 in the water maze during training is evaluated by the time spent (mean ± SEM, top) and distance travelled (mean ± SEM, bottom) before climbing onto the platform (one-way ANOVA for repeated measures, time: F(6) = 3.685, p = 0.014, η^2^ = 0.551; distance: F(6) = 3.604, p = 0.016, η^2^ = 0.546). **D**, The success rate on each training day increased monotonically before spatial probe tests. **E**, The performance of tree shrews in the water maze during training. Both escape latency (top) and escape distance (bottom) plateaued after five days. Data from individual animals are shown in color lines, while the mean ± SEM of data from twelve animals is represented by the black trace. **F**, TS095 in the first spatial probe test. The trajectory of the animal’s head is indicated by black traces and the location of the platform is represented by a dark circle. The occupancy in the target quadrant in distance is indicated by the number at bottom left. This animal crossed the platform location tree times. **G-I**, The number of platform (PF) crosses (**G**), percentage of time (**H**) and travel distance (**I**) in the target quadrant (TQ) of each animal in the two spatial probe tests. The same color code is used as in panel **E**. n.s., not significant.

In the spatial probe test (probe 1), the trajectory of each animal was monitored for the first 60 seconds. The tree shrews were expected to search around and cross the platform location a couple of times (**Fig. 4 F**). To evaluate their performance in the probe test, we used two types of measures that were commonly used in rodents: the number of platform location crossings (a conservative measure), and their occupancy time and distance in the target quadrant (sensitive measures) (Maei et al., 2009). Group data analysis showed that the tree shrews crossed the platform 2.42 ± 0.34 times (**Fig. 4 G**), and the occupancy time and distance in the target quadrant were only slightly above a nonbiased search pattern (equal distribution of 0.25) (time: 0.304 ± 0.020 one-sample t-test, n = 12, t = 2.690, p = 0.021; distance: 0.295 ± 0.018, t = 2.528, p = 0.028) (**Fig. 4 H-I**). We attributed this to insufficient training before the probe test, thus extended the swimming-pool task by three more days of training without changing the platform location (days 8-10), followed by a second spatial probe test (probe 2). The improvement in spatial learning during excess training was limited: a trend towards statistical significance was observed in both escape latency and distance when comparing days 8-10 with days 5-7 (two-way ANOVA for repeated measures, n = 12, latency: F(1,2) = 4.579, p = 0.056, η^2^ = 0.294; distance: F(1,2) = 3.677, p = 0.081, η^2^ = 0.251) (**Fig. 4 D-E**). Furthermore, the recall of platform location in the second probe test, evaluated by number of platform crossings (3.00 ± 0.54), was not better than the first probe test (Wilcoxon signed rank test, Z = 10.21, p = 0.307) (**Fig. 4 G**), suggesting that excessive training did not significantly improve memory recall of the platform location in the swimming pool. Interestingly, tree shrews’ target quadrant occupancy in probe test 2 dropped to equal distribution of 0.25 (time: 0.259 ± 0.025; distance: 0.257 ± 0.022; one-sample t-test, both p values ≥ 0.915), although statistically not different from probe 1 (one-way ANOVA for repeated measures, n = 12, time: F(1) = 3.890, p = 0.074, η^2^ = 0.261; distance: F(1) = 3.263, p = 0.098, η^2^ = 0.229) (**Fig. 4 H-I**), suggesting that the target quadrant occupancy in probe tests might not be a reliable measure to evaluate tree shrew’s memory of platform location. Overall, the study concluded that excessive training did not lead to significant improvements in spatial memory acquisition or retrieval in the swimming pool and that a seven-day training is sufficient for tree shrew to learn the water maze test.

### 3.4. Various lesion sizes in the hippocampus

In order to compare the sensitivity to changes in spatial memory across different paradigms, we performed hippocampal lesioning on twelve tree shrews that had been tested in all three mazes (**Fig. 1 A**). Colchicine infusions were made to lesion the hippocampal formation, including the dentate gyrus, hippocampus proper, and subiculum (Keuker et al., 2003), along the dorsoventral axis in both hemispheres. We aimed to infuse the dorsal, intermediate, and ventral portions of the hippocampus on each side, with priority given to the dorsal hippocampi which are known to be more involved in spatial memory (Moser et al., 1993; Moser et al., 1995; Moser and Moser, 1998). The actual number of infusions depended on the state of the animal during surgery. We were able to make six infusion sites as planned in six tree shrews, whereas five animals received 3-5 injections due to difficulties encountered during surgery (**Tab. 1**). One animal (TS104) died during surgery, possibly due to isoflurane overdose, thus was excluded from further analyses.

**Table 1.** Extent of the hippocampal lesion in each tree shrew.

| Animal | Left hippocampus |  |  | Right hippocampus |  |  | Total | Mean |
| --- | --- | --- | --- | --- | --- | --- | --- | --- |
|  | Inject | Volume | Lesion | Inject | Volume | Lesion | volume | lesion |
|  | sites | (mm<sup>3</sup>) | size (%) | sites | (mm<sup>3</sup>) | size (%) | (mm<sup>3</sup>) | size (%) |
| TS094 | 3 | 15.56 | 80.08 | 3 | 12.70 | 83.74 | 28.25 | 81.91 |
| TS095 | 3 | 6.66 | 91.47 | 3 | 3.01 | 96.15 | 9.67 | 93.81 |
| TS096 | 2 | 24.18 | 69.04 | 2 | 29.78 | 61.87 | 53.96 | 65.46 |
| TS097 | 1 | 54.30 | 30.47 | 2 | 31.03 | 60.27 | 85.33 | 45.37 |
| TS099 | 3 | 6.98 | 91.06 | 3 | 10.97 | 85.95 | 17.95 | 88.51 |
| TS100 | 1 | 59.11 | 24.31 | 2 | 32.32 | 58.61 | 91.43 | 41.46 |
| TS101 | 3 | 6.08 | 92.22 | 3 | 5.74 | 92.65 | 11.82 | 92.43 |
| TS102 | 3 | 4.07 | 94.79 | 3 | 3.61 | 95.38 | 7.68 | 95.08 |
| TS103 | 2 | 34.06 | 56.39 | 2 | 36.48 | 53.28 | 70.54 | 54.84 |
| TS106 | 3 | 8.62 | 88.97 | 3 | 25.49 | 67.36 | 34.11 | 78.16 |
| TS108 | 3 | 6.67 | 91.46 | 2 | 25.77 | 67.00 | 32.44 | 79.23 |

After completing cognitive tests, histology was obtained to assess the extent of the hippocampal lesions in the remaining animals. The border between damaged and healthy hippocampal tissue was sharp thus can be clearly defined (**Fig. 5 A**). The volume of healthy hippocampal tissue was quantified for each animal and the proportion of lesion was estimated using a standard tree shrew hippocampus with a volume of 78.10 mm^3^ (**Fig. 5 B**). The volume of healthy tissue varied between animals, ranging from 7.68 mm^3^ to 91.43 mm^3^, which corresponded to lesions sizes between 41.46% and 95.08% (**Tab. 1**). Lesion size was not quantified in each subregion, as the tri-synaptic circuit involving all structures within a transverse block is required for hippocampal function (Stepan et al., 2015). Minor inadvertent damage was observed in the para-hippocampal region, including the entorhinal cortex, pre- and para-subiculum, of TS095 and TS102, whose hippocampi were most effectively lesioned. An unintended cortical lesion was also observed in TS106, possibly caused by unsuccessful injections to the right hippocampus, which damaged the piriform cortex but spared part of the dorsal dentate gyrus (**Extended Fig. 5**).

**Figure 5.**
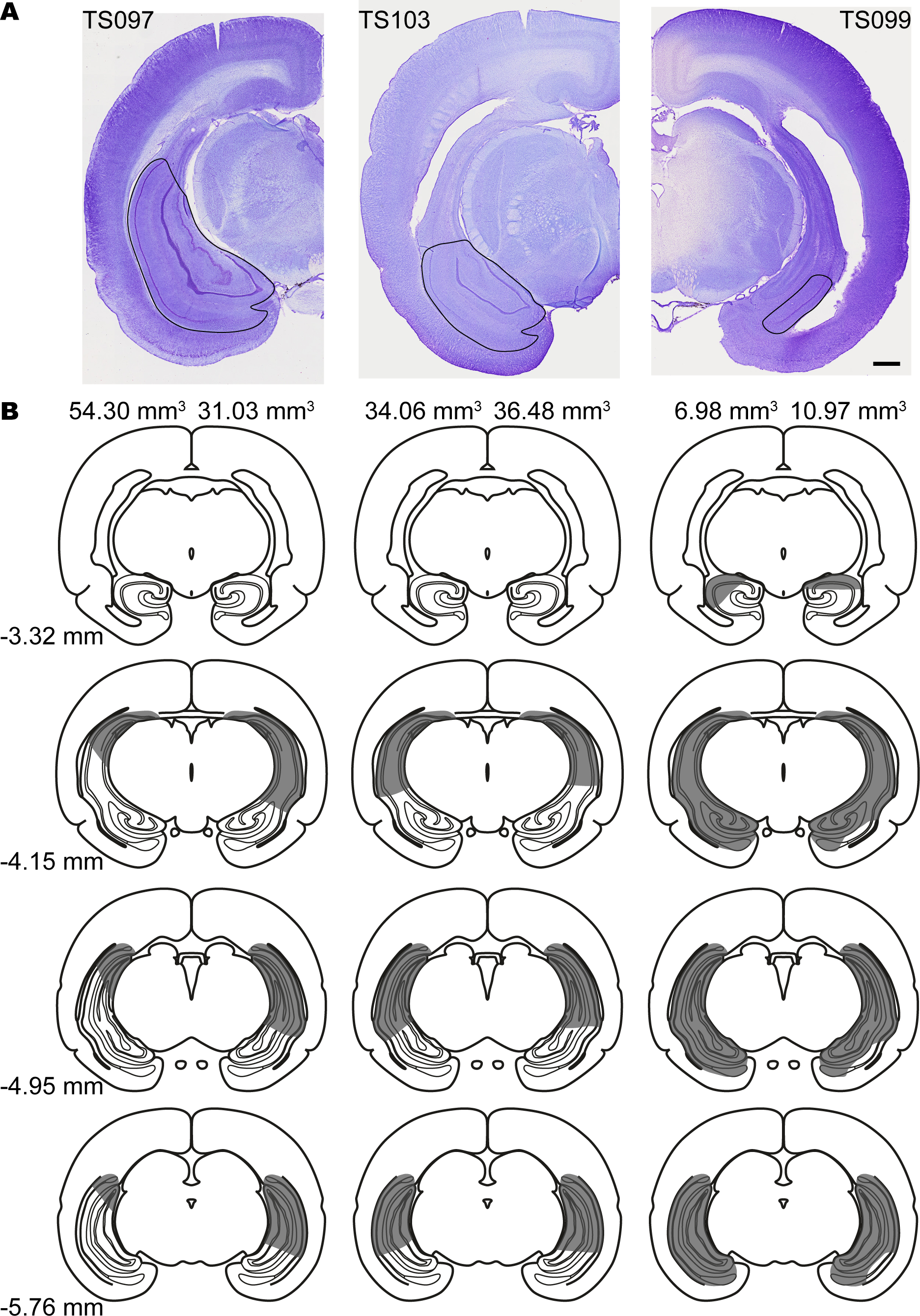
Representative lesions in the hippocampus. **A**. Brain sections from three animals with different levels of hippocampal lesion. The healthy hippocampal tissue is encircled by black traces. Scale bar: 1 mm. **B**, Representative schematics of equally spaced brain sections illustrating various degrees of hippocampal lesion in TS097 (left), TS103 (middle) and TS099 (right). Shadings indicate lesions in the hippocampus. The numbers on top of the schematics indicate volumes of healthy hippocampus in each hemisphere. The numbers at bottom left refer to the distance posterior to Bregma. The original schematics used in this figure were adapted from *The tree shrew (Tupaia belangeri chinensis) brain in stereotaxic coordinates* (Zhou and Ni, 2016).

### 3.5. Degenerated reference memory in the radial-arm task after hippocampal lesion

Eleven tree shrews were tested in the radial-arm maze again at least three weeks after lesions to both hippocampi (**Fig. 1 A**). Arms 3, 6, and 8 were baited after lesion, which had a symmetrical spatial arrangement to the baited arms in the previous test. The animals were tested for seven consecutive days, with similar numbers of trials performed as in the previous test. Ten out of eleven animals performed well in the task, except TS106, who entered the arms randomly every day without consuming the rewards. Unlike the other animals, TS106’s working memory and reference memory error rates remained high over the test days after the lesion (one-way ANOVA for repeated measures for TS106, 25 trials, working memory: F(6) = 1.158, p = 0.332, η^2^ = 0.046; reference memory: F(6) = 1.623, p = 0.145, η^2^ = 0.063; other animals: all p values ≤ 0.002). We attributed this poor performance to TS106’s sudden weight gain during the recovery period after the lesion, which compromised its motivation to perform the task (**Fig. 1 G**). TS106 was removed from further analysis in the radial-arm maze. The frequency of the favorite route and route switches when visiting baited arms did not change significantly after the lesion (Test days 1-7, preferred sequence: pre-lesion: 0.599 ± 0.036, post-lesion: 0.561 ± 0.066, paired t-test, t(9) = 0.493, p = 0.634; sequence switches: pre-lesion: 0.473 ± 0.082, post-lesion: 0.481 ± 0.159, paired t-test, t(9) = 0.113, p = 0.912), indicating that tree shrews completed both rounds of radial-arm tasks using the same strategy, in the first seven days.

We first compared tree shrew’s working memory on pre-and post-lesion test days 1-7, and found it was not impaired after the lesion (**Fig. 6 A-B**, left). Our analysis of group data indicated that working memory performance significantly improved over the test days after hippocampal lesion, similar to pre-lesion trials (two-way ANOVA for repeated measures, 10 animals, F(6) = 30.775, p < 0.001, η^2^ = 0.774). We did not observe any significant difference in working memory performance before and after the lesion (F(1) = 2.895, p = 0.123, η^2^ = 0.243), nor did we find any interaction between hippocampal lesion and test days (F(1,6) = 1.782, p = 0.120, η^2^ = 0.165) (**Fig. 6 C**, left). Although it has long been debated whether the hippocampus is involved in working memory, our findings were consistent with a recent report that systematically analyzed twenty-six human studies and concluded that working memory does not activate the hippocampus (Slotnick, 2022). In contrast to working memory, reference memory remained unchanged in tree shrews with relatively small hippocampal damage, but was impaired when both hippocampi were mostly destroyed. (**Fig. 6 A-B**, right). Our analysis of group data indicated that tree shrews’ reference memory after the lesion also improved over test days, but was significantly impaired (two-way ANOVA for repeated measures, 10 animals, test day: F(6) = 45.159, p < 0.001, η^2^ = 0.834; lesion: F(1) = 7.717, p = 0.021, η^2^ = 0.462), with maximum change on day 6 (post-hoc analysis with Bonferroni correction). Again, we did not observe any lesion × test day interaction (F(1,6) = 1.090, p = 0.380, η^2^ = 0.108) (**Fig. 6 C**, right). These findings suggest that the hippocampus plays a critical role in reference memory, but not in working memory, when tree shrews performing the radial-arm task.

**Figure 6.**
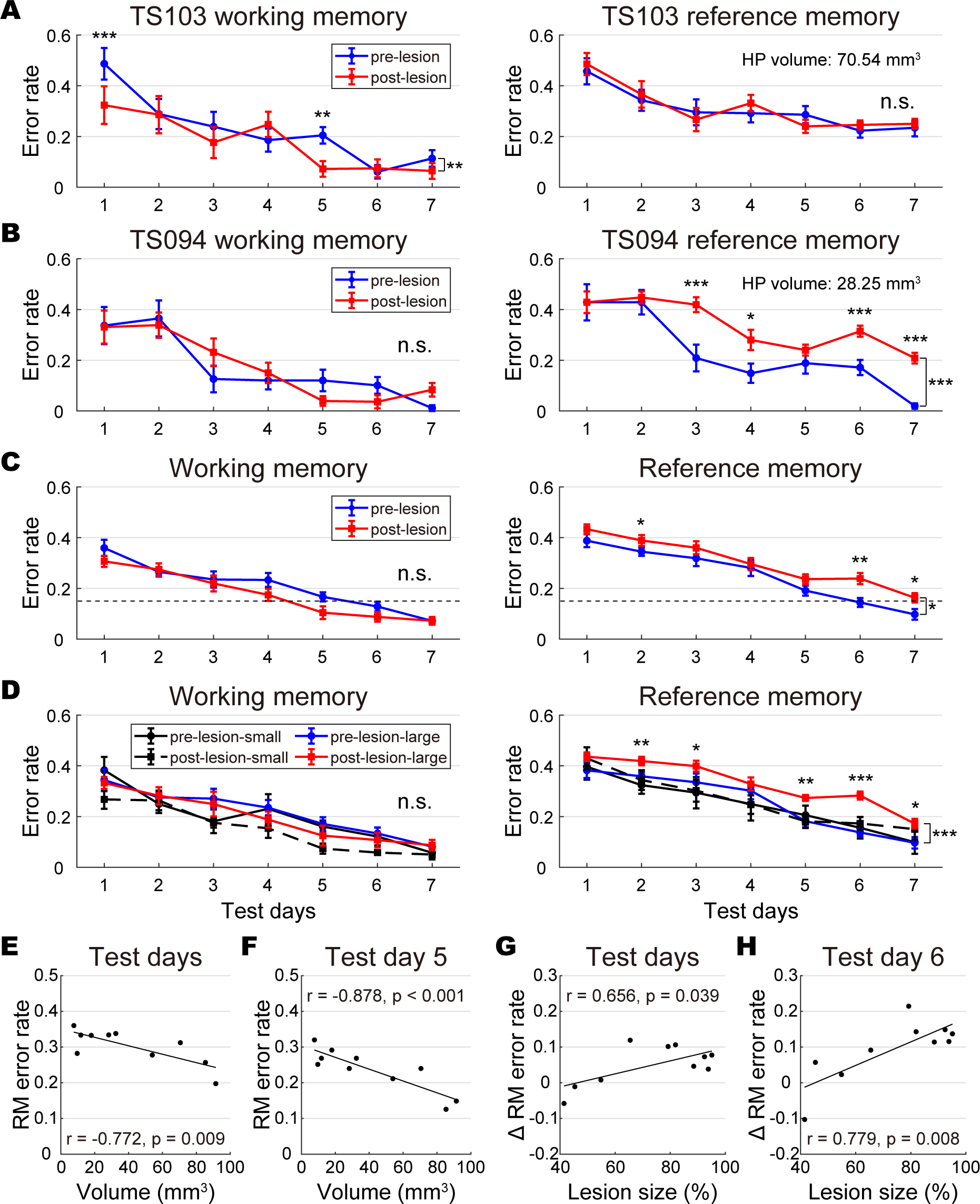
Performance in the radial-arm maze after hippocampal lesion. **A**-**B**, Working memory (mean ± SEM, left) and reference memory (mean ± SEM, right) performance of tree shrews with small (**A**) or large hippocampal lesions (**B**). TS103 exhibited a slight increase in working memory performance on test days 1 and 5 (two-way ANOVA for repeated measures, test day: F(6) = 18.095, p < 0.001, η^2^ = 0.430; lesion: F(1) = 8.218, p = 0.008, η^2^ = 0.255; lesion × test day: F(1,6) = 3.678, p = 0.002, η^2^ = 0.133), while reference memory remained unchanged (test day: F(6) = 17.539, p < 0.001, η^2^ = 0.422; lesion: F(1) = 0.227, p = 0.638, η^2^ = 0.009; lesion × test day: F(1,6) = 0.874, p = 0.516, η^2^ = 0.035). TS094’s working memory was not altered (test day: F(6) = 27.555, p < 0.001, η^2^ = 0.534; lesion: F(1) = 0.136, p = 0.716, η^2^ = 0.006; lesion × test day: F(1,6) = 3.040, p = 0.008, η^2^ = 0.112), while reference memory was impaired after the lesion (test day: F(6) = 45.539, p < 0.001, η^2^ =0.655; lesion: F(1) = 51.936, p < 0.001, η^2^ = 0.684; lesion × test day: F(1,6) = 5.410, p < 0.001, η^2^ = 0.184). **C**, Working memory (left) and reference memory (right) of tree shrews included in the analysis, before and after the lesion (mean ± SEM). Dashed lines indicate the 0.15 threshold for task learning. Note that the reference memory error rates were consistently higher than 0.15 after lesion. **D**, Task performance in the radial-ram maze of animals grouped by volume of healthy hippocampal tissue (mean ± SEM). **E**, Scatter plots showing significant correlations between volume of healthy hippocampal tissue and reference memory (RM) error rates averaged across seven test days after the lesion. The regression line is shown in black. **F**, Same as in panel **E**, but showing data for post-lesion test day 5, when the correlation was the most prominent. **G**, Scatter plots showing significant correlation between the extent of the hippocampal lesion and changes in reference memory over the course of seven test days after the lesion. **H**, same as in panel **G**, but showing data on post-lesion test day 6, when the difference in reference memory was most poonounced. n.s., not significant, *, p < 0.05, **, p < 0.01, ***, p < 0.001.

To further clarify the relationship between hippocampal function and reference memory, we divided the tree shrews into two groups based on the size of their healthy hippocampal tissue. The first group had a small hippocampal lesion (volume > 50 mm^3^), while the second group had a large lesion (volume < 50 mm^3^). We found that both groups had unchanged working memory after the lesion, as demonstrated by two-way ANOVA for repeated measures (small-lesion group: 4 animals, test day: F(6) = 12.618, p < 0.001, η^2^ = 0.808; lesion: F(1) = 2.209, p = 0.234, η^2^ = 0.424; lesion × test day: F(1,6) = 2.221, p = 0.089, η^2^ = 0.425; large-lesion group, 6 animals, test day: F(6) = 16.648, p < 0.001, η^2^ = 0.769; lesion: F(1) = 0.752, p = 0.425, η^2^ = 0.131; lesion × test day: F(1,6) = 0.528, p = 0.782, η^2^ = 0.096) (**Fig. 6 D**, left). Visits to un-baited arms were not affected in the small-lesion group after hippocampal lesion (two-way ANOVA for repeated measures, 4 animals, test day: F(6) = 12.890, p < 0.001, η^2^ = 0.811; lesion: F(1) = 0.153, p = 0.722, η^2^ = 0.049; lesion × test day: F(1,6) = 0.291, p = 0.933, η^2^ = 0.089), but increased significantly in the large-lesion group (two-way ANOVA for repeated measures, 6 animals, test day: F(6) = 34.871, p < 0.001, η^2^ = 0.875; lesion: F(1) = 42.217, p = 0.001, η^2^ = 0.894; lesion × test day: F(1,6) = 2.119, p = 0.080, η^2^ = 0.298), on five out of seven test days (post-hoc analysis with Bonferroni correction).

We found a strong relationship between size of healthy tissue in the hippocampus and reference memory in tree shrew, evidenced by significant correlations between volume of remaining tissue and reference memory error rates averaged across post-lesion test days (**Fig. 6 E**). Our analysis of everyday data revealed that this volume-performance correlation was also significant on test days 3-6 (Pearson’s correlation, p values ≤ 0.03) (**Fig. 6 F**). Additionally, we investigated whether the extent of hippocampal damage affected spatial memory, and found that the reduction in reference memory was indeed associated with lesion size (**Fig. 6 G**). This correlation was strongest on test day 6, when the largest difference in spatial memory was observed (**Fig. 6 H**). Taken together, our findings indicated that tree shrews’ reference memory in the radial-arm maze was impaired by significant hippocampal lesions, while working memory remained largely unaffected. Furthermore, the declines in reference memory were correlated with the size of the lesion in both hippocampi. These conclusions were also supported by analyses that include data from TS106 (data not shown).

### 3.6. Dramatic declines in post-lesion spatial learning in the cheeseboard maze

The hippocampus-lesioned tree shrews were next re-tested in the cheeseboard maze (**Fig. 1 A**). To ensure consistency between the two rounds of tests, we baited the wells in a manner similar to the pre-lesion test (see methods). Although group data from the previous test plateaued from day 3 (**Fig. 3 F**), performance of individual animal fluctuated over days. Therefore, we tested the animals consecutively for 5 days. Similar to previous report in rats (Olton and Werz, 1978), the tree shrews’ running speed in the maze was higher after hippocampal lesion, especially on early test days (**Fig. 1 H**), suggesting that their locomotion abilities were not impaired by the lesion. Moreover, the animals employed the same strategy to complete the tasks, as evidenced by the consistent degree of route preference in both rounds of tests (two-way ANOVA for repeated measures, 11 animals, frequency of favorite sequence: test day: F(4) = 1.884, p = 0.132, η^2^ = 0.159; lesion: F(1) = 0.426, p = 0.529, η^2^ = 0.041; lesion × test day: F(1,4) = 1.021, p = 0.408, η^2^ = 0.093; frequency of sequence switches: test day: F(4) = 1.005, p = 0.417, η^2^ = 0.091; lesion: F(1) = 0.010, p = 0.922, η^2^ = 0.001; lesion × test day: F(1,4) = 0.391, p = 0.814, η^2^ = 0.038) (**Fig. 7 A-B**).

**Figure 7.**
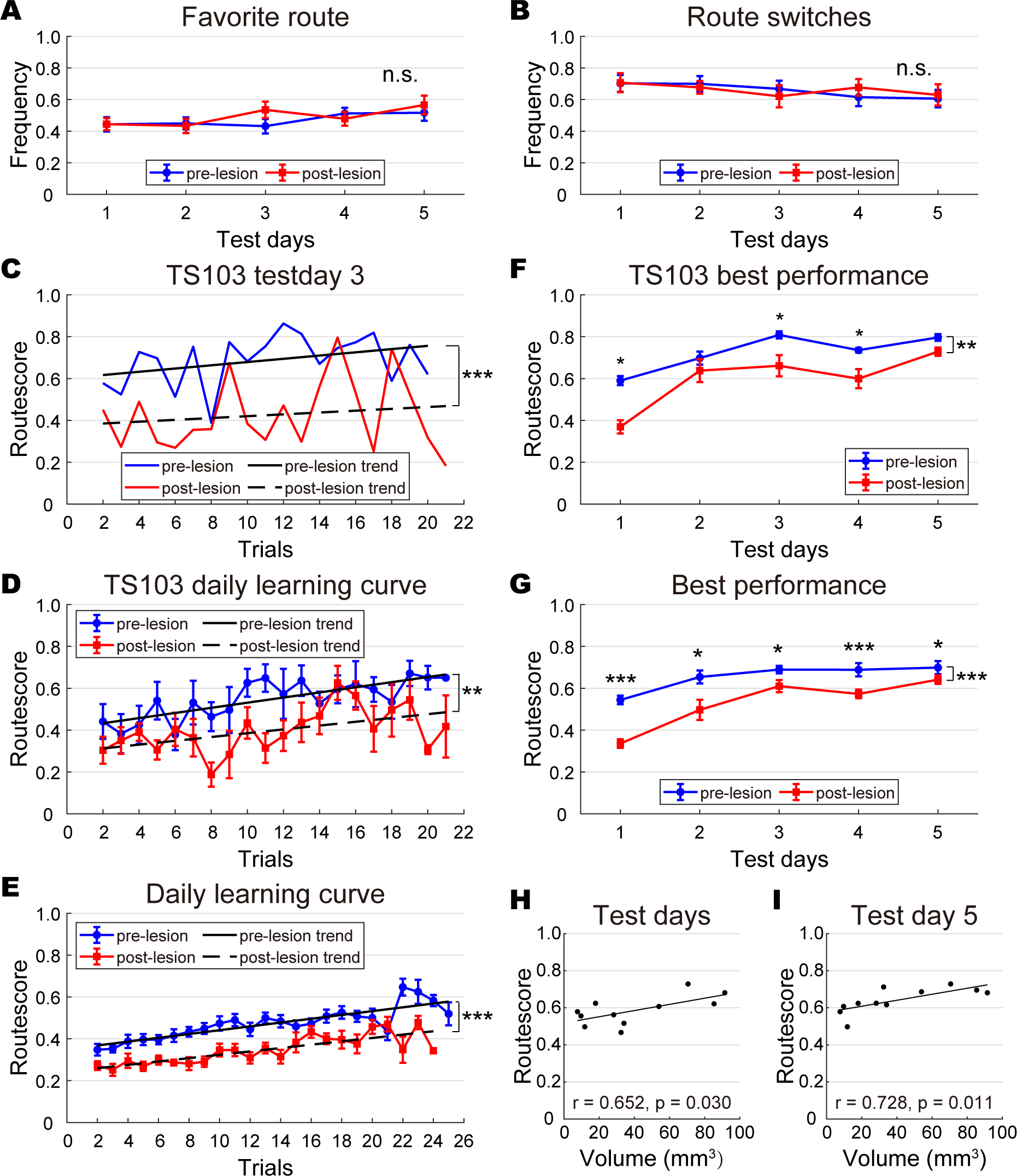
Performance in the cheeseboard maze after hippocampal lesion. **A-B**, Tree shrews’ route preference on pre- and post-lesion days, measured by frequency of favorite route (mean ± SEM, **A**) and frequency of route switches (mean ± SEM, **B**). **C**, TS103’s learning curve on test day 3, before and after the lesion (paired t-test, t(19) = 6.053, p < 0.001). **D**, TS103’s daily learning curve (mean ± SEM), averaged across five test days (two-way ANOVA for repeated measures, trial: F(19) = 2.781, p = 0.001, η^2^ = 0.410; lesion: F(1) = 22.307, p = 0.009, η^2^ = 0.848; lesion × trial: F(1,19) = 1.337, p = 0.186, η^2^ = 0.251). **E**, The daily learning curve (mean ± SEM) of tree shrews in the cheeseboard maze, averaged across five test days. **F**, TS103’s best performance (mean ± SEM) before and after the lesion, averaged from five best trials on each test day (two-way ANOVA for repeated measures, test day: F(4) = 19.192, p < 0.001, η^2^ = 0.828; lesion: F(1) = 38.068, p = 0.004, η^2^= 0.905; lesion × test day: F(1,4) = 2.188, p = 0.117, η^2^ = 0.354). **G**, Same as in panel **F**, the best performance (mean ± SEM) on each test day averaged across all animals. **H**, Scatter plots showing a significant correlation between the volume of healthy hippocampal tissue and the best performance averaged across five test days after the lesion. The regression line is shown in black. **I**, Same as in panel **H**, but showing the correlation between residual hippocampal volume and the best performance on post-lesion test day 5. n.s., not significant, *, p < 0.05, **, p < 0.01, ***, p < 0.001.

The animals were able to partially remember the locations of reward wells after undergoing more than 20 trials of training, but their routes to the rewards were always longer than those on the same test day prior to the lesion (**Fig. 7 C**). Combining test days also revealed a significant reduction in routescores (**Fig. 7 D**). Analysis of learning curves from all animals revealed that their memory of reward locations in the cheeseboard maze gradually improved within 25 trials in both rounds of tests (two-way ANOVA for repeated measures, 11 animals, F(22) = 13.291, p < 0.001, η^2^ = 0.571), but was significantly impaired after bilateral hippocampal lesion (F(1) = 64.935, p < 0.001, η^2^ = 0.867). A significant trial × lesion interaction was also observed (F(1,22) = 3.684, p < 0.001, η^2^ = 0.269) (**Fig. 7 E**). The best performance was significantly reduced after lesion for every individual tree shrew (two-way ANOVA for repeated measures, all p values ≤ 0.036) (**Fig. 7 F**). Our analysis of group data also revealed a significant reduction in best performance (two-way ANOVA for repeated measures, 11 animals, test day: F(4) = 59.011, p < 0.001, η^2^ = 0.855; lesion: F(1) = 44.780, p < 0.001, η^2^= 0.817; lesion × test day: F(1,4) = 3.759, p = 0.011, η^2^ = 0.273), and the reduction was significant on every test day (post-hoc analysis with Bonferroni correction) (**Fig. 7 G**).

We next asked whether post-lesion task performance was affected by hippocampal lesion size. Our analysis revealed a significant correlation between the volume of healthy hippocampal tissue and the averaged routescores (**Fig. 7 H**), and the correlation was most pronounced on test day 5 (**Fig. 7 I**). In conclusion, these results indicated that tree shrews’ spatial learning in the cheeseboard task was dramatically impaired by bilateral hippocampal lesions, even in animals with the smallest damages, raising the possibility that this paradigm is sensitive to subtle changes in spatial memory.

### 3.7. Impaired spatial memory retrieval in the water maze test after lesion

The tree shrews were subjected to a final test in the water maze (**Fig. 1 A**). We moved the platform from quadrant 2 to quadrant 3 for this round of test. To our surprise, unlike the other two tasks, the tree shrews behaved much better after lesion and their performance in the swimming pool on post-lesion days 1-4 was similar to pre-lesion days 4-7. Under such circumstances, it was deemed unnecessary to compare spatial learning (memory encoding) in the water maze before and after the lesion. Therefore, we modified the protocol by testing the tree shrews after 4 days of training (probe 1), when they had displayed comparable escaping behavior. Following the first probe test, the animals were trained for an additional three days without relocating the platform (days 5-7), and finally their memory retrieval was tested for a second time (probe 2), which was similar to the previous experiment.

For the sake of convenience, post-lesion training days 1-7 were aligned with pre-lesion training days 4-10 in the analyses. The individual learning curves of animals after lesion did not differ significantly from those during pre-lesion training in the two training stages. (two-way ANOVA for repeated measures, all p values ≥ 0.139) (**Fig. 8 A**). Group data indicated that the success rates in training trials were close to being statistically significant after lesion (paired t-test, first stage: t(3) = 2.784, p = 0.069; second stage: t(2) = 4.150, p = 0.053). Both escape latency and distance decreased over time during the first stage of training (two-way ANOVA for repeated measures, 11 animals, latency: F(3) = 6.164, p = 0.002, η^2^ = 0.381; distance: F(3) = 3.316, p = 0.040, η^2^ = 0.239), but not in the second stage (latency: F(2) = 1.943, p = 0.169, η^2^ = 0.163; distance: F(2) = 1.922, p = 0.172, η^2^ = 0.161). There was no difference observed between pre- and post-lesion training days in escape latency (first stage: F(1) = 1.100, p = 0.319, η^2^ = 0.099; second stage: F(1) = 2.601, p = 0.138, η^2^ = 0.206) or escape distance (first stage: F(1) = 1.122, p = 0.314, η^2^ = 0.101; second stage: F(1) = 1.820, p = 0.207, η^2^ = 0.154). No lesion × training day interaction was detected in these tests (first stage: latency: F(1,3) = 1.668, p = 0.195, η^2^ = 0.143; distance: F(1,3) = 1.493, p = 0.237, η^2^ = 0.130; second stage: latency: F(1,2) = 0.964, p = 0.398, η^2^ = 0.088; distance: F(1,2) = 1.075, p = 0.360, η^2^ = 0.097) (**Fig. 8 B**). These results indicated that spatial memory acquisition was comparable before retrieval tests, suggesting that even a small piece of remaining hippocampal tissue can support spatial learning in the water maze. Our findings in tree shrew were consistent with previous studies in rat (Moser et al., 1993; Moser et al., 1995; Moser and Moser, 1998).

**Figure 8.**
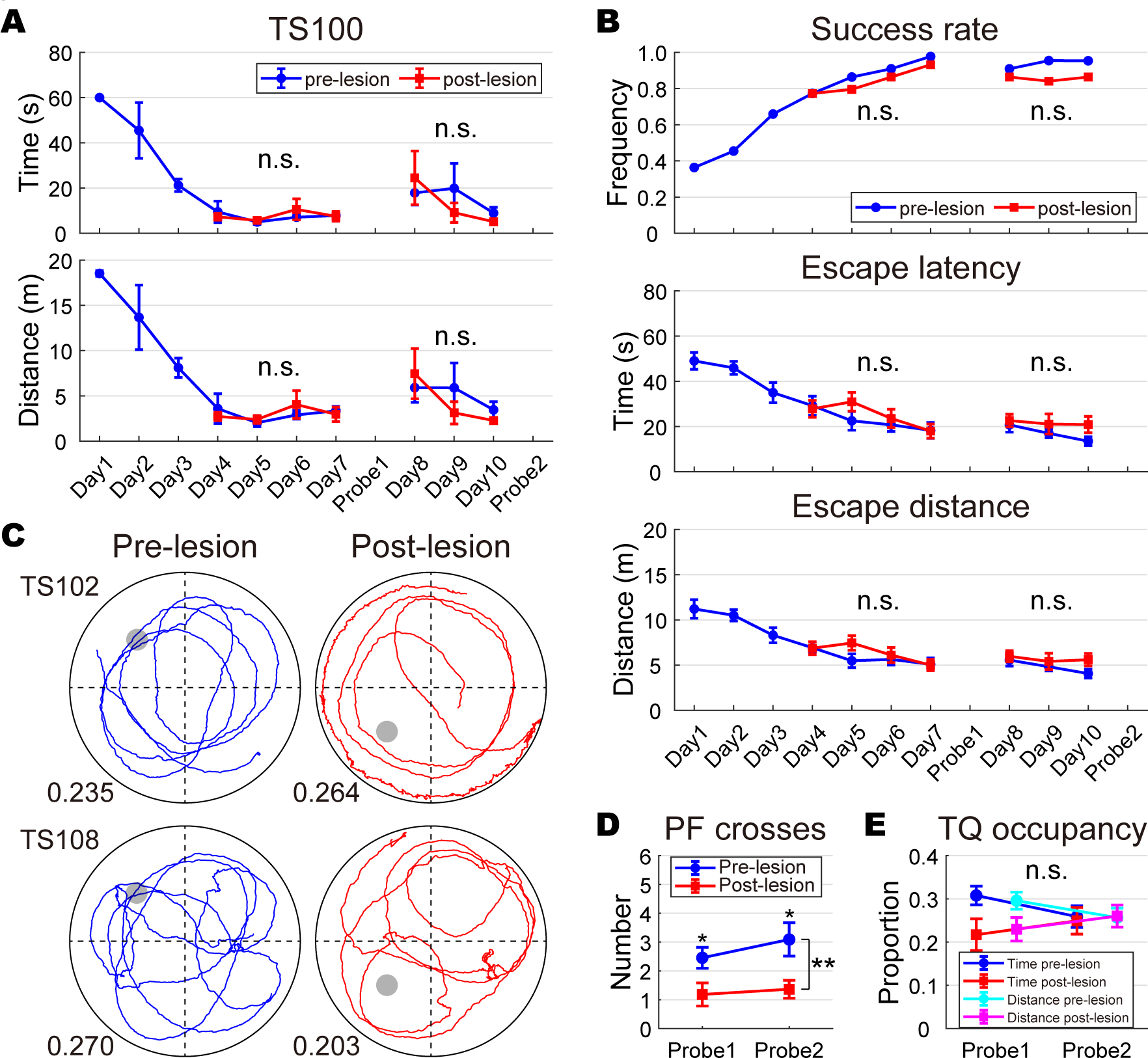
Performance in the water maze after hippocampal lesion. **A**. TS100’s performance on training days before and after the lesion, measured by escape latency (mean ± SEM, top) and distance (mean ± SEM, bottom). Post-lesion days 4-10 in the plots were training days 1-7 after lesion. Both time and distance were comparable after realignment (two-way ANOVA for repeated measures, all p values ≥ 0.232). **B**, The success rate (top), escape latency (mean ± SEM, middle) and distance (mean ± SEM, bottom) on pre- and post-lesion training days. **C**, The trajectories of TS102 and TS108 in probe test 1, before and after hippocampal lesion, indicated by blue and red traces, respectively. The location of the platform is represented by a dark circle. The target quadrant occupancy in distance is indicated at bottom left of each trajectory plot. **D-E**, The number of platform (PF) crosses (mean ± SEM, **D**) and the target quadrant (TQ) occupancy (mean ± SEM, **E**) in spatial probe tests before and after the lesion. n.s., not significant, *, p < 0.05, **, p < 0.01.

We next compared tree shrews’ retrieval of spatial memory in the probe tests. The number of platform crossing was reduced in some animals after lesion, while the target quadrant occupancy remained largely the same (**Fig. 8 C**). Our analysis of group data showed that the numbers of platform crossings were similar between the first and second probe tests (post-lesion test 1: 1.18 ± 0.40; test 2: 1.36 ± 0.31; two-way ANOVA for repeated measures, 11 animals, F(1) = 1.000, p = 0.341, η^2^ = 0.091), but were significantly reduced after lesion (F(1) = 12.692, p = 0.005, η^2^ = 0.559) in both tests (post-hoc analysis with Bonferroni correction) (**Fig. 8 D**). No lesion × test interaction was detected (F(1,1) = 0.387, p = 0.548, η^2^= 0.037). Tree shrews’ target quadrant occupancy in both probe tests after lesion were again very close to equal distribution of 0.25 (one-sample t-test, both p values ≥ 0.392). No difference was observed in occupancy after lesion (time: lesion: F(1) = 3.246, p = 0.102, η^2^ = 0.245; test: F(1) = 0.142, p = 0.714, η^2^ = 0.014; lesion × test: F(1,1) = 1.393, p = 0.265, η^2^ = 0.122; distance: lesion: F(1) = 2.233, p = 0.166, η^2^= 0.183; test: F(1) = 0.046, p = 0.834, η^2^ = 0.005; lesion × test: F(1,1) = 1.510, p = 0.247, η^2^ = 0.131) (**Fig. 8 E**), suggesting that tree shrews may employ a different strategy than rodents when searching for the platform. Different from the other two tasks, no correlation was observed between hippocampal lesion size and spatial learning during training, or recall of platform location in probe tests (Pearson correlation, 11 animals, all p values ≥ 0.115). To conclude, our results suggested that recall of the platform location in the swimming pool was impaired after hippocampal lesion, even when comparable levels of spatial learning had been achieved. This implies that there is a dissociation between encoding and retrieval of spatial memory in the hippocampus, which is consistent with previous reports both in rat and human (Zeineh et al., 2003; Lee and Kesner, 2004; Eldridge et al., 2005; Duncan et al., 2014).

## 4. Discussion

Tree shrew is a viable animal model for research on AD (Fan et al., 2018), a neurodegenerative disease impairs spatial cognition in its early stage (Coughlan et al., 2018). Current choices of cognitive paradigms evaluating spatial memory in tree shrew are limited. In this study, we developed paradigms to assess spatial memory in tree shrew, and compared their robustness by testing a group of twelve animals before and after bilateral lesions to the hippocampus. We found that hippocampal lesion compromised task performance in all mazes, but the degree of deterioration varied across paradigms. Our results suggest that the cheeseboard task is the most appropriate choice among the three spatial paradigms when evaluating spatial memory impairments in tree shrew, which can potentially help to monitor progressive cognitive declines in aging or disease models.

### 4.1. Landmark-based navigation in the spatial tasks

Behavioral paradigms testing spatial memory typically require animals to navigate in the mazes based on distal landmarks (visuospatial navigation), rather than local cues (beacon navigation) (Nyberg et al., 2022). To achieve this, we tested the animals in a well-lit room enriched with distal visual cues, removed all possible visual and tactile marks near the rewards, and obscured the rewards’ smell to minimize local cues (see methods). Tree shrews initially struggled to follow the odor cues but soon learned to ignore them during pre-training. This observation suggested that tree shrews had learnt to use distal visual cues instead of proximal order cues to locate the rewards. This was further supported by observations from another group of four animals in our lab. We tested them in a cue-deprived version of radial-arm maze, where distal landmarks were mostly not visible. Those tree shrews were unable to locate the baited arms after three weeks of training, but instead developed a kinesthetic strategy to visit adjacent arms consecutively (data not shown). To sum up, in the current study, tree shrews used the distal landmarks to position themselves in the mazes, suggesting that their internal navigation system was involved when performing the tasks.

### 4.2. Varied spatial memory demand across behavior paradigms

In the present study, we compared the spatial memory of tree shrews before and after hippocampal lesion, to avoid potential inter-individual behavioral differences. This experimental design allowed each animal to serve as its own control and enabled us to compare the three paradigms in the same animal. The spatial learning in the cheeseboard maze and the radial-arm maze was compromised after the lesion (**Fig. 6 C**, **Fig. 7 E&G**), while the memory retrieval was impaired in the water maze test (**Fig. 8 C-D**). Moreover, compromised spatial learning was observed in the cheeseboard task but not in the radial-arm task in tree shrews with relatively small hippocampal damage (**Fig. 6 A**, **Fig. 7 C, D&F**). Overall, our findings suggested that the cheeseboard maze was the most sensitive to loss of spatial memory among the three paradigms. Importantly, the minimal impact of previous experiences in the cheeseboard maze and its reservoir of reward locations, allowed for multiple repetitions of the task in the same animals, which is critical for monitoring progressive changes in spatial memory during aging, and the development of AD.

The differential effects of hippocampal damage on these paradigms may be attributed to varying levels of spatial coding in the entorhinal-hippocampal circuit. Specifically, we observed that animals were required to memorize three target locations in both the cheeseboard and radial-arm mazes, whereas only one platform location was necessary in the water maze. This difference implied that more hippocampal neurons were recruited to represent multiple goal locations in the reward-based paradigms (Dupret et al., 2010; Sarel et al., 2017; Spiers et al., 2018). In addition, we noticed that the cheeseboard paradigm required the animals to update reward locations on a daily basis. Daily memory of reward positions in the cheeseboard maze could only be stabilized by short inter-trial replays in the hippocampus, whereas spatial memory initially encoded in the hippocampus, could be enhanced and transmitted to the neocortex by post-training consolidation during sleep in the other two tasks, where target locations remained stationary (Klinzing et al., 2019; Brodt et al., 2023). Therefore, hippocampal lesion may be partially compensated by memory consolidation in the radial-arm task and the swimming-pool task, but not in the cheeseboard task. Finally, we found that animals in the cheeseboard maze had to compute both distance and direction to locate the reward positions, whereas in the radial-arm maze only direction was necessary to identify the baited arms. Studies have shown that the processing of distance and direction coding occurs independently through separate populations of neurons located in distinct layers of the medial entorhinal cortex (Fyhn et al., 2004; Hafting et al., 2005; Sargolini et al., 2006), which project both locally, and to different subregions of the hippocampus (Canto et al., 2008). Based on our observations, we suggest that the convergence of distance and direction information is more likely to occur in the hippocampus rather than the entorhinal cortex, as the integrity of the hippocampus was critical for completing the cheeseboard task.

### 4.3. Interspecies variability in the water maze

In this study, we found that tree shrews were able to learn the platform location faster in the second round of water maze test, even after bilateral lesions to the hippocampus (**Fig. 8 A-B**). This suggests that previous experience strongly influences spatial learning on repeated tests in the water maze. In addition, our own observations indicated that tree shrews may be less adept at swimming compared to rats, which could partially explain why more practice is required before they can effectively navigate by distal landmarks when performing water-based tasks (Whishaw and Tomie, 1996). To further support this hypothesis, we compared the performance of eleven male Long-Evans rats of similar age (22 weeks) using the same swimming pool and training protocol (**Fig. 9**). The results showed that rats learned the platform location significantly faster than tree shrews. However, our results from dry-land tasks in this study suggested that the impairments displayed by the same group of tree shrews in the swimming pool were not due to inadequacy in spatial memory *per se*, but rather due to other non-spatial limiting factors such as swimming abilities. The variations in swimming skills may also lead to different searching strategies in the water maze. It was observed that tree shrews’ occupancy in the target quadrant during probe tests was close to a no-biased search pattern (**Fig. 4 H-I**, **Fig. 8 C&E**), which differed from previous reports in rats (Morris, 1984; Moser et al., 1993; Moser et al., 1995; Moser and Moser, 1998). We speculated that tree shrew may use a more flexible platform-searching strategy than rodents during probe tests. Overall, our findings suggested that water-based cognitive tests may introduce confounds in tree shrew, whereas behavior paradigms in dry-land mazes are more likely to produce optimal results when assessing spatial memory in this species. The present study also provided insights into the comparative cognitive abilities of different species and highlights the importance of considering species-specific factors when designing and interpreting cognitive experiments.

**Figure 9.**
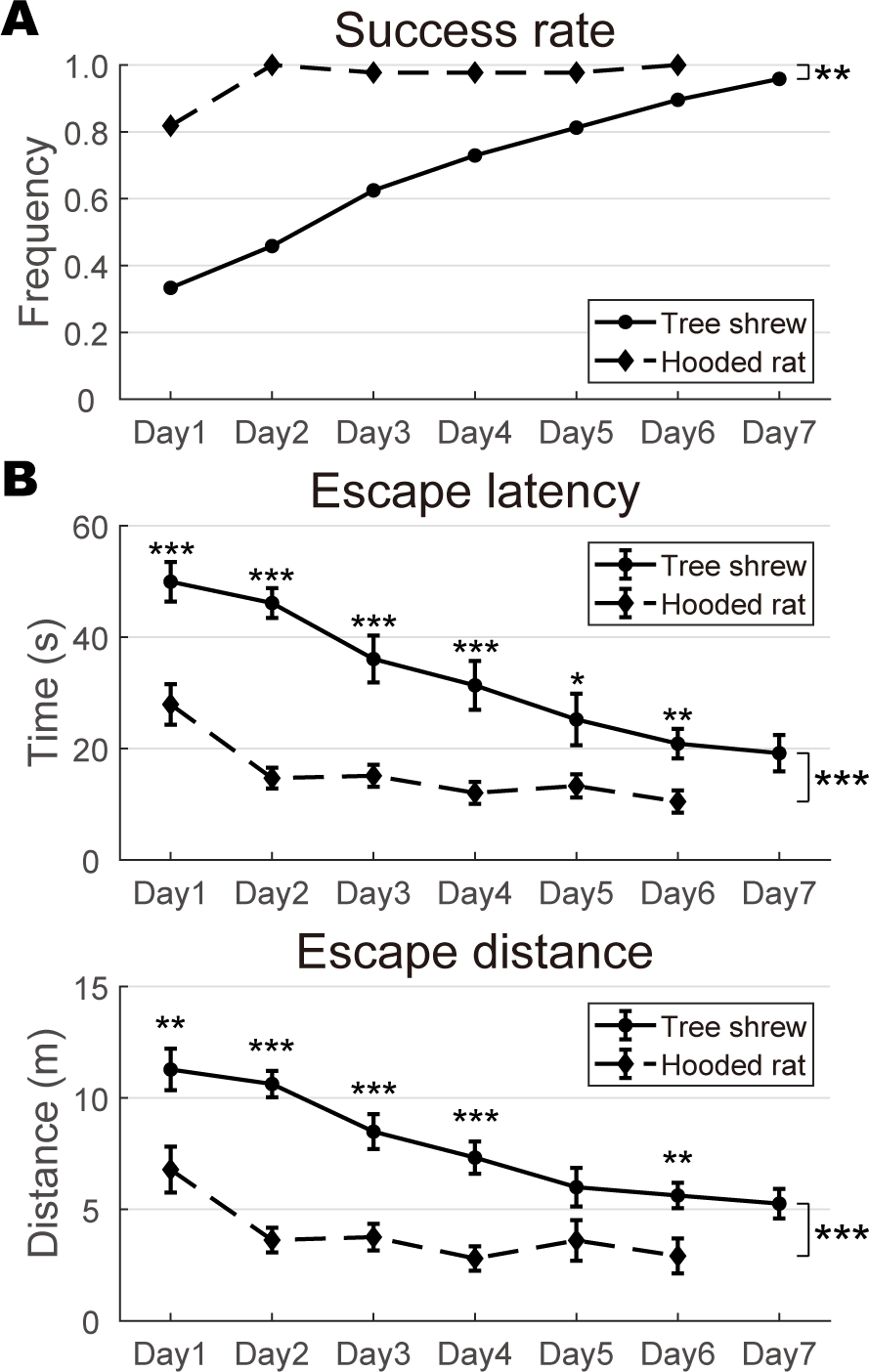
Fast learning of hooded rats in the water maze. **A**, The success rates of rats and tree shrews on each training day. Rats had significantly higher success rates than tree shrews (paired t-test, t(5) = 4.423, p = 0.007). **B**, The learning curves (mean ± SEM) in the water maze during training for both species, the rats found the platform significantly faster than the tree shrews, reflected both by escape latency (top) and escape distance (bottom) (two-way ANOVA for repeated measures, 23 animals, escape latency: species: F(1) = 41.544, p < 0.001, η^2^ = 0.664; training day: F(5) = 20.489, p < 0.001, η^2^= 0.494; training day × species: F(1,5) = 4.207, p = 0.002, η^2^ = 0.167; escape distance: species: F(1) = 40.543, p < 0.001, η^2^ = 0.659; training day: F(5) = 15.136, p < 0.001, η^2^ = 0.419; training day × species: F(1,5) = 3.301, p = 0.008, η^2^ = 0.136).

### 4.4. Potential application in spatial cognitive studies

Tree shrew possesses phylogenetic proximity to primates (Fan et al., 2013; Fan et al., 2019), reflected not only by its closely resembled genome and gene expression patten (Xu et al., 2012; Fan et al., 2013; Fan et al., 2018), but also by similar brain morphology (Remple et al., 2006; Remple et al., 2007; Wong and Kaas, 2009; Wang et al., 2013; Ni et al., 2016; Dai et al., 2017; Ni et al., 2018) and comparable higher order cognitive functions (Mustafar et al., 2018; Jiang et al., 2021; Pan et al., 2022). The tree shrew has advantages as a model animal in cognitive studies of spatial navigation (Savier et al., 2021), due to its well-developed visual system (MacEvoy et al., 2009; Lee et al., 2016; Petry and Bickford, 2019; Tanabe et al., 2022), prefrontal cortex (Parra et al., 2019), and possibly internal navigation system (Finkelstein et al., 2016), in addition to disease models (Cao et al., 2003; Xiao et al., 2017; Yao, 2017; Li et al., 2018). Numerous studies have revealed how the entorhinal-hippocampal circuit represents the cognitive map of the external world (Moser et al., 2015; Rowland et al., 2016). However, it remains unclear how animals use this cognitive map to flexibly navigate to spatial goals (Long and Lu, 2022; Nyberg et al., 2022). In addition, both traditional and cutting-edge approaches monitoring neural activities in the brain, including in vivo electrophysiology (Dimanico et al., 2021) and two-photon calcium imaging (Lee et al., 2016; Schumacher et al., 2022) are now being transferred from rodents to tree shrew. Collectively, these lines of evidence have raised the possibility that the tree shrew is an ideal and viable animal for cognitive studies related to flexible navigation with multiple goals and dynamic goal-directed routes. The cheeseboard maze, which has the potential to provide enormous combinations of reward sites and massive choices of routes, is particularly suitable for flexible spatial tasks.

## Supporting information

Extended data for Figure 5

## Acknowledgements

This study was supported by Science and Technology Innovation (STI) 2030-Major Projects 2022ZD0205000 and Startup from Kunming Institute of Zoology, Chinese Academy of Sciences. The authors would like to thank Ya-Li Duan, Shu-Yi Hu, Xun Tang and Yu-Ming Sun for technical assistance in hippocampal infusion and DeepLabCut; Chu Deng, Xiao-Fan Ge, Jia-Li Long, Xiao-Nan Zhao and Yi-Fan Ye for handling the animals; Hai-Bing Xu, Dun Mao, Yong Gu and Yong-Gang Yao for suggestions to behavior paradigms and the manuscript.

## Author contributions

L.L., C.J.L., and Y.Q.H. designed the experiment. C.J.L. and Y.Q.H. performed tree shrew behavior tests. H.Y.Z. and X.C. performed the rat water maze test. C.J.L. and R.Z. made lesions. C.J.L. and Y.Q.H. quantified the hippocampal lesion, analyzed the task performance, and made figures. L.L. supervised the project and wrote the manuscript. All authors contributed to discussion and interpretation.

## Code Accessibility

All custom MATLAB scripts used in this study are available upon request.

## References

Adams JN, Kim S, Rizvi B, Sathishkumar M, Taylor L, Harris AL, Mikhail A, Keator DB, McMillan L, Yassa MA (2022) Entorhinal-Hippocampal Circuit Integrity Is Related to Mnemonic Discrimination and Amyloid-P Pathology in Older Adults. Journal of Neuroscience 42:8742–8753.

Braak H, Braak E (1991) Neuropathological stageing of Alzheimer-related changes. Acta Neuropathol 82:239–259.

Brodt S, Inostroza M, Niethard N, Born J (2023) Sleep-A brain-state serving systems memory consolidation. Neuron 111:1050–1075.

Canto CB, Wouterlood FG, Witter MP (2008) What does the anatomical organization of the entorhinal cortex tell us? Neural Plast 2008:381243.

Cao J, Yang EB, Su JJ, Li Y, Chow P (2003) The tree shrews: adjuncts and alternatives to primates as models for biomedical research. J Med Primatol 32:123–130.

Chen ZY, Zhang Y (2022) Animal models of Alzheimer’s disease: Applications, evaluation, and perspectives. Zool Res 43:1026–1040.

Coughlan G, Laczo J, Hort J, Minihane AM, Hornberger M (2018) Spatial navigation deficits - overlooked cognitive marker for preclinical Alzheimer disease? Nat Rev Neurol 14:496–506.

Dai JK, Wang SX, Shan D, Niu HC, Lei H (2017) A diffusion tensor imaging atlas of white matter in tree shrew. Brain Struct Funct 222:1733–1751.

Dimanico MM, Klaassen AL, Wang J, Kaeser M, Harvey M, Rasch B, Rainer G (2021) Aspects of tree shrew consolidated sleep structure resemble human sleep. Commun Biol 4:722.

Dubois B et al. (2014) Advancing research diagnostic criteria for Alzheimer’s disease: the IWG-2 criteria. Lancet Neurology 13:614–629.

Duncan K, Tompary A, Davachi L (2014) Associative encoding and retrieval are predicted by functional connectivity in distinct hippocampal area CA1 pathways. J Neurosci 34:11188–11198.

Dupret D, O’Neill J, Pleydell-Bouverie B, Csicsvari J (2010) The reorganization and reactivation of hippocampal maps predict spatial memory performance. Nat Neurosci 13:995–1002.

Eldridge LL, Engel SA, Zeineh MM, Bookheimer SY, Knowlton BJ (2005) A dissociation of encoding and retrieval processes in the human hippocampus. J Neurosci 25:3280–3286.

Fan Y, Luo R, Su LY, Xiang Q, Yu D, Xu L, Chen JQ, Bi R, Wu DD, Zheng P, Yao YG (2018) Does the genetic feature of the Chinese tree shrew (Tupaia belangeri chinensis) support Its potential as a viable model for Alzheimer’s disease research? J Alzheimers Dis 61:1015–1028.

Fan Y, Ye MS, Zhang JY, Xu L, Yu DD, Gu TL, Yao YL, Chen JQ, Lv LB, Zheng P, Wu DD, Zhang GJ, Yao YG (2019) Chromosomal level assembly and population sequencing of the Chinese tree shrew genome. Zool Res 40:506–521.

Fan Y et al. (2013) Genome of the Chinese tree shrew. Nat Commun 4:1426.

Finkelstein A, Las L, Ulanovsky N (2016) 3-D maps and compasses in the brain. Annu Rev Neurosci 39:171–196.

Fu H, Rodriguez GA, Herman M, Emrani S, Nahmani E, Barrett G, Figueroa HY, Goldberg E, Hussaini SA, Duff KE (2017) Tau Pathology Induces Excitatory Neuron Loss, Grid Cell Dysfunction, and Spatial Memory Deficits Reminiscent of Early Alzheimer’s Disease. Neuron 93:533–541 e535.

Fuchs E (2005) Social stress in tree shrews as an animal model of depression: an example of a behavioral model of a CNS disorder. CNS Spectr 10:182–190.

Fyhn M, Molden S, Witter MP, Moser EI, Moser MB (2004) Spatial representation in the entorhinal cortex. Science 305:1258–1264.

Hafting T, Fyhn M, Molden S, Moser MB, Moser EI (2005) Microstructure of a spatial map in the entorhinal cortex. Nature 436:801–806.

Igarashi KM, Lu L, Colgin LL, Moser MB, Moser EI (2014) Coordination of entorhinal-hippocampal ensemble activity during associative learning. Nature 510:143–147.

Jiang M, Wang M, Shi Q, Wei L, Lin Y, Wu D, Liu B, Nie X, Qiao H, Xu L, Yang T, Wang Z (2021) Evolution and neural representation of mammalian cooperative behavior. Cell Rep 37:110029.

Jun H, Bramian A, Soma S, Saito T, Saido TC, Igarashi KM (2020) Disrupted Place Cell Remapping and Impaired Grid Cells in a Knockin Model of Alzheimer’s Disease. Neuron 107:1095–1112 e1096.

Keuker JI, Rochford CD, Witter MP, Fuchs E (2003) A cytoarchitectonic study of the hippocampal formation of the tree shrew (Tupaia belangeri). J Chem Neuroanat 26:1–15.

Khani A, Rainer G (2012) Recognition memory in tree shrew (Tupaia belangeri) after repeated familiarization sessions. Behav Processes 90:364–371.

Klinzing JG, Niethard N, Born J (2019) Mechanisms of systems memory consolidation during sleep. Nat Neurosci 22:1598–1610.

Lee I, Kesner RP (2004) Encoding versus retrieval of spatial memory: Double dissociation between the dentate gyrus and the perforant path inputs into CA3 in the dorsal hippocampus. Hippocampus 14:66–76.

Lee KS, Huang X, Fitzpatrick D (2016) Topology of ON and OFF inputs in visual cortex enables an invariant columnar architecture. Nature 533:90–94.

Li R, Zanin M, Xia X, Yang Z (2018) The tree shrew as a model for infectious diseases research. J Thorac Dis 10:S2272–S2279.

Li Z, He X, Wu S, Huang R, Li H, Wang Z, Wang L, Qin D, Kong Y, Guo Y, Ma X, Turck CW, Xiong Z, Wang W, Hu X (2022) Naturally occurring Alzheimer’s disease in rhesus monkeys. bioRxiv:2022.2010.2020.513120.

Lin N, Xiong LL, Zhang RP, Zheng H, Wang L, Qian ZY, Zhang P, Chen ZW, Gao FB, Wang TH (2016) Injection of Abeta1-40 into hippocampus induced cognitive lesion associated with neuronal apoptosis and multiple gene expressions in the tree shrew. Apoptosis 21:621–640.

Long JL, Lu L (2022) Dynamic coding in the hippocampus during navigation. Zool Res 43:1023–1025.

MacEvoy SP, Tucker TR, Fitzpatrick D (2009) A precise form of divisive suppression supports population coding in the primary visual cortex. Nat Neurosci 12:637–645.

Maei HR, Zaslavsky K, Teixeira CM, Frankland PW (2009) What is the Most Sensitive Measure of Water Maze Probe Test Performance? Front Integr Neurosci 3:4.

Masters CL, Bateman R, Blennow K, Rowe CC, Sperling RA, Cummings JL (2015) Alzheimer’s disease. Nat Rev Dis Primers 1:15056.

Masuda Y, Odashima J, Murai S, Saito H, Itoh M, Itoh T (1994) Radial arm maze behavior in mice when a return to the home cage serves as the reinforcer. Physiol Behav 56:785–788.

Mathis A, Mamidanna P, Cury KM, Abe T, Murthy VN, Mathis MW, Bethge M (2018) DeepLabCut: markerless pose estimation of user-defined body parts with deep learning. Nat Neurosci 21:1281–1289.

Morris R (1984) Developments of a water-maze procedure for studying spatial learning in the rat. J Neurosci Methods 11:47–60.

Moser EI, Moser MB, Andersen P (1993) Spatial learning impairment parallels the magnitude of dorsal hippocampal lesions, but is hardly present following ventral lesions. J Neurosci 13:3916–3925.

Moser MB, Moser EI (1998) Distributed encoding and retrieval of spatial memory in the hippocampus. J Neurosci 18:7535–7542.

Moser MB, Rowland DC, Moser EI (2015) Place cells, grid cells, and memory. Cold Spring Harbor perspectives in biology 7:a021808.

Moser MB, Moser EI, Forrest E, Andersen P, Morris RG (1995) Spatial learning with a minislab in the dorsal hippocampus. Proc Natl Acad Sci U S A 92:9697–9701.

Mustafar F, Harvey MA, Khani A, Arato J, Rainer G (2018) Divergent solutions to visual problem solving across mammalian species. eNeuro 5.

Nair J, Topka M, Khani A, Isenschmid M, Rainer G (2014) Tree shrews (Tupaia belangeri) exhibit novelty preference in the novel location memory task with 24-h retention periods. Front Psychol 5:303.

Ni RJ, Luo PH, Shu YM, Chen JT, Zhou JN (2016) Whole-brain mapping of afferent projections to the bed nucleus of the stria terminalis in tree shrews. Neuroscience 333:162–180.

Ni RJ, Huang ZH, Luo PH, Ma XH, Li T, Zhou JN (2018) The tree shrew cerebellum atlas: Systematic nomenclature, neurochemical characterization, and afferent projections. J Comp Neurol 526:2744–2775.

Ni RJ, Tian Y, Dai XY, Zhao LS, Wei JX, Zhou JN, Ma XH, Li T (2020) Social avoidance behavior in male tree shrews and prosocial behavior in male mice toward unfamiliar conspecifics in the laboratory. Zool Res 41:258–272.

Nyberg N, Duvelle E, Barry C, Spiers HJ (2022) Spatial goal coding in the hippocampal formation. Neuron 110:394–422.

Ohl F, Oitzl MS, Fuchs E (1998) Assessing cognitive functions in tree shrews: visuo-spatial and spatial learning in the home cage. J Neurosci Methods 81:35–40.

Olton DS, Samuelson RJ (1976) Remembrance of Places Passed - Spatial Memory in Rats. J Exp Psychol-Anim B 2:97–116.

Olton DS, Werz MA (1978) Hippocampal function and behavior: spatial discrimination and response inhibition. Physiol Behav 20:597–605.

Pan TT, Liu C, Li DM, Nie BB, Zhang TH, Zhang W, Zhao SL, Zhou QX, Liu H, Zhu GH, Xu L, Shan BC (2022) Nucleus accumbens-linked executive control networks mediating reversal learning in tree shrew brain. Zool Res 43:528–531.

Parra A, Baker CA, Bolton MM (2019) Regional Specialization of Pyramidal Neuron Morphology and Physiology in the Tree Shrew Neocortex. Cereb Cortex 29:4488–4505.

Paspalas CD, Carlyle BC, Leslie S, Preuss TM, Crimins JL, Huttner AJ, van Dyck CH, Rosene DL, Nairn AC, Arnsten AFT (2018) The aged rhesus macaque manifests Braak stage III/IV Alzheimer’s-like pathology. Alzheimers Dement 14:680–691.

Pawlik M, Fuchs E, Walker LC, Levy E (1999) Primate-like amyloid-beta sequence but no cerebral amyloidosis in aged tree shrews. Neurobiol Aging 20:47–51.

Petry HM, Bickford ME (2019) The Second Visual System of The Tree Shrew. J Comp Neurol 527:679–693.

Pryce CR, Fuchs E (2017) Chronic psychosocial stressors in adulthood: Studies in mice, rats and tree shrews. Neurobiol Stress 6:94–103.

Remple MS, Reed JL, Stepniewska I, Kaas JH (2006) Organization of frontoparietal cortex in the tree shrew (Tupaia belangeri). I. Architecture, microelectrode maps, and corticospinal connections. J Comp Neurol 497:133–154.

Remple MS, Reed JL, Stepniewska I, Lyon DC, Kaas JH (2007) The organization of frontoparietal cortex in the tree shrew (Tupaia belangeri): II. Connectional evidence for a frontal-posterior parietal network. J Comp Neurol 501:121–149.

Rowland DC, Roudi Y, Moser MB, Moser EI (2016) Ten Years of Grid Cells. Annu Rev Neurosci 39:19–40.

Sarel A, Finkelstein A, Las L, Ulanovsky N (2017) Vectorial representation of spatial goals in the hippocampus of bats. Science 355:176–180.

Sargolini F, Fyhn M, Hafting T, McNaughton BL, Witter MP, Moser MB, Moser EI (2006) Conjunctive representation of position, direction, and velocity in entorhinal cortex. Science 312:758–762.

Savier E, Sedigh-Sarvestani M, Wimmer R, Fitzpatrick D (2021) A bright future for the tree shrew in neuroscience research: Summary from the inaugural Tree Shrew Users Meeting. Zool Res 42:478–481.

Schneider CA, Rasband WS, Eliceiri KW (2012) NIH Image to ImageJ: 25 years of image analysis. Nat Methods 9:671–675.

Schumacher JW, McCann MK, Maximov KJ, Fitzpatrick D (2022) Selective enhancement of neural coding in V1 underlies fine-discrimination learning in tree shrew. Curr Biol 32:3245–3260 e3245.

Shang S, Wang C, Guo C, Huang X, Wang L, Zhang C (2015) The formation and extinction of fear memory in tree shrews. Front Behav Neurosci 9:204.

Slotnick SD (2022) Does working memory activate the hippocampus during the late delay period? Cogn Neurosci-Uk 13:182–207.

Spiers HJ, Olafsdottir HF, Lever C (2018) Hippocampal CA1 activity correlated with the distance to the goal and navigation performance. Hippocampus 28:644–658.

Squire LR (1992) Memory and the hippocampus: a synthesis from findings with rats, monkeys, and humans. Psychol Rev 99:195–231.

Steffenach HA, Witter M, Moser MB, Moser EI (2005) Spatial memory in the rat requires the dorsolateral band of the entorhinal cortex. Neuron 45:301–313.

Stepan J, Dine J, Eder M (2015) Functional optical probing of the hippocampal trisynaptic circuit in vitro: network dynamics, filter properties, and polysynaptic induction of CA1 LTP. Front Neurosci 9:160.

Tanabe S, Fu J, Cang J (2022) Strong tuning for stereoscopic depth indicates orientation-specific recurrent circuitry in tree shrew V1. Curr Biol 32:5274–5284 e5276.

Vorhees CV, Williams MT (2006) Morris water maze: procedures for assessing spatial and related forms of learning and memory. Nat Protoc 1:848–858.

Wang L, Lu J, Zeng Y, Guo Y, Wu C, Zhao H, Zheng H, Jiao J (2020) Improving Alzheimer’s disease by altering gut microbiota in tree shrews with ginsenoside Rg1. FEMS Microbiol Lett 367.

Wang S, Shan D, Dai J, Niu H, Ma Y, Lin F, Lei H (2013) Anatomical MRI templates of tree shrew brain for volumetric analysis and voxel-based morphometry. J Neurosci Methods 220:9–17.

Whishaw IQ, Tomie J (1996) Of mice and mazes: similarities between mice and rats on dry land but not water mazes. Physiol Behav 60:1191–1197.

Wong P, Kaas JH (2009) Architectonic subdivisions of neocortex in the tree shrew (Tupaia belangeri). Anat Rec (Hoboken) 292:994–1027.

Xiao J, Liu R, Chen CS (2017) Tree shrew (Tupaia belangeri) as a novel laboratory disease animal model. Zool Res 38:127–137.

Xu L, Chen SY, Nie WH, Jiang XL, Yao YG (2012) Evaluating the phylogenetic position of Chinese tree shrew (Tupaia belangeri chinensis) based on complete mitochondrial genome: implication for using tree shrew as an alternative experimental animal to primates in biomedical research. J Genet Genomics 39:131–137.

Yamashita A, Fuchs E, Taira M, Hayashi M (2010) Amyloid beta (Abeta) protein- and amyloid precursor protein (APP)-immunoreactive structures in the brains of aged tree shrews. Curr Aging Sci 3:230–238.

Yamashita A, Fuchs E, Taira M, Yamamoto T, Hayashi M (2012) Somatostatin-immunoreactive senile plaque-like structures in the frontal cortex and nucleus accumbens of aged tree shrews and Japanese macaques. J Med Primatol 41:147–157.

Yao YG (2017) Creating animal models, why not use the Chinese tree shrew (Tupaia belangeri chinensis)? Zool Res 38:118–126.

Ye MS, Zhang JY, Yu DD, Xu M, Xu L, Lv LB, Zhu QY, Fan Y, Yao YG (2021) Comprehensive annotation of the Chinese tree shrew genome by large-scale RNA sequencing and long-read isoform sequencing. Zool Res 42:692-709.

Ying J, Keinath AT, Lavoie R, Vigneault E, El Mestikawy S, Brandon MP (2022) Disruption of the grid cell network in a mouse model of early Alzheimer’s disease. Nat Commun 13:886.

Zambello E, Fuchs E, Abumaria N, Rygula R, Domenici E, Caberlotto L (2010) Chronic psychosocial stress alters NPY system: different effects in rat and tree shrew. Prog Neuropsychopharmacol Biol Psychiatry 34:122–130.

Zeineh MM, Engel SA, Thompson PM, Bookheimer SY (2003) Dynamics of the hippocampus during encoding and retrieval of face-name pairs. Science 299:577–580.

Zheng YT, Yao YG, Xu L (2014) Basic biology and disease models of tree shrews. Kunming: Yunnan Science and Technology Press.

Zhou J-N, Ni R-J (2016) The tree shrew (Tupaia belangeri chinensis) brain in stereotaxic coordinates. In, p 1 online resource (592 p. Singapore: Springer,.

