## Extended data for Figure 5 for "A comparison of behavior paradigms assessing spatial memory in tree shrew"

Figure 5-1

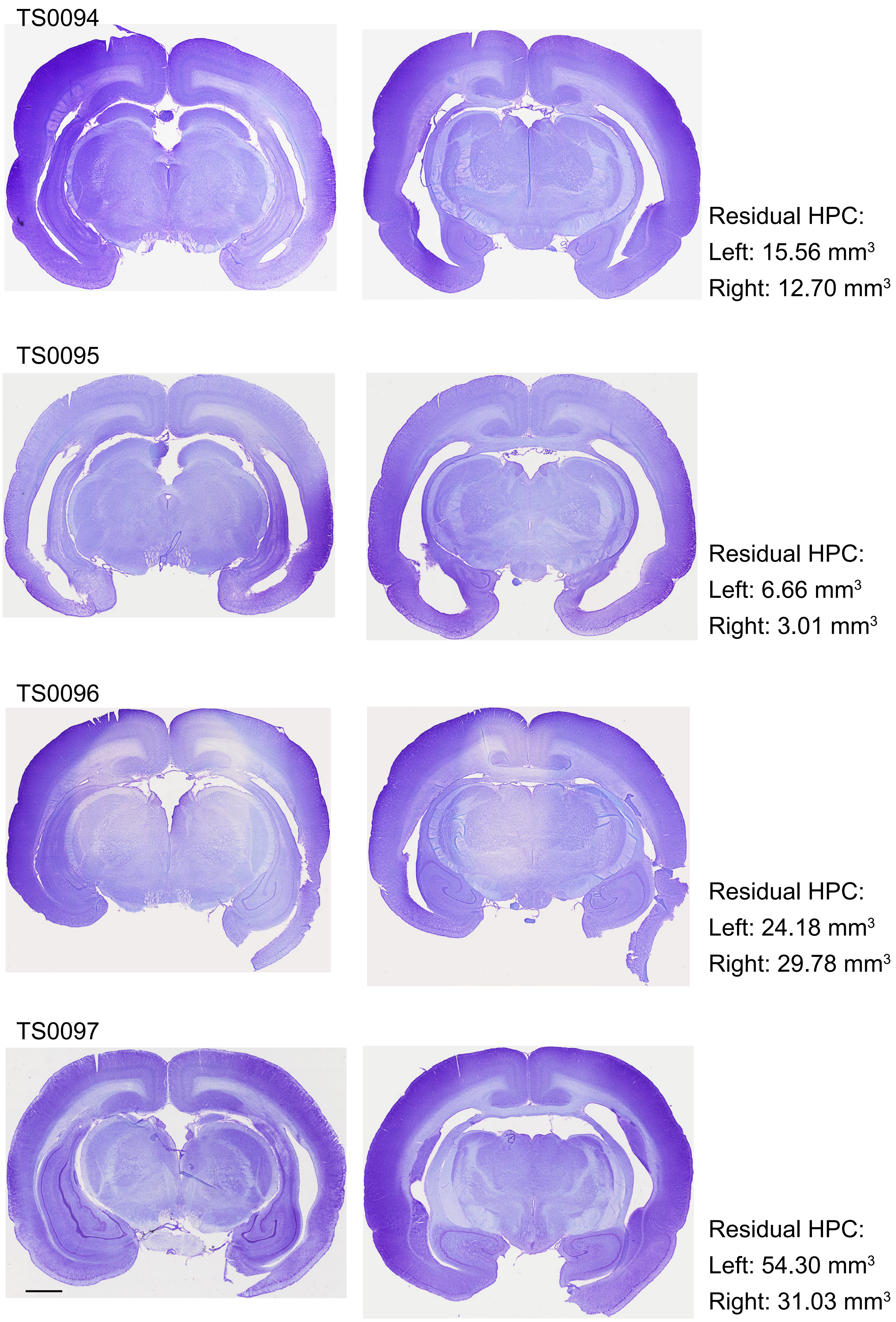

TS0094
Residual HPC:
Left: 15.56 mm³
Right: 12.70 mm³
TS0095
Residual HPC:
Left: 6.66 mm³
Right: 3.01 mm³
TS0096
Residual HPC:
Left: 24.18 mm³
Right: 29.78 mm³
TS0097
Residual HPC:
Left: 54.30 mm³
Right: 31.03 mm³

Figure 5-2

TS0099

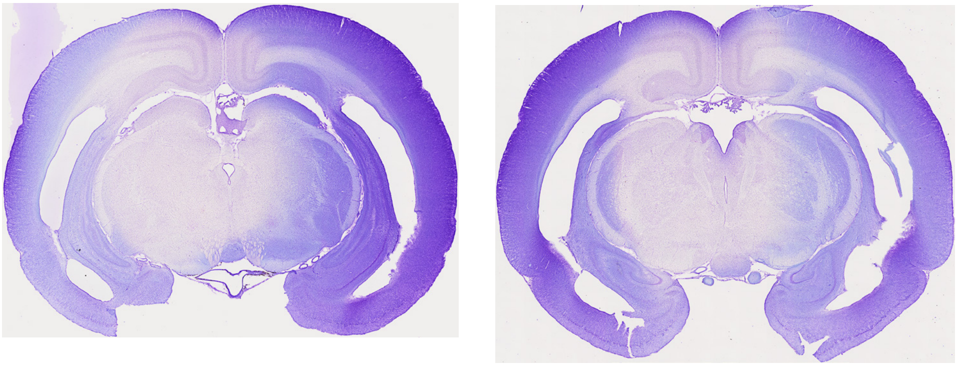

Residual HPC:
Left: 6.98 $mm^3$
Right: 10.97 $mm^3$

TS0100

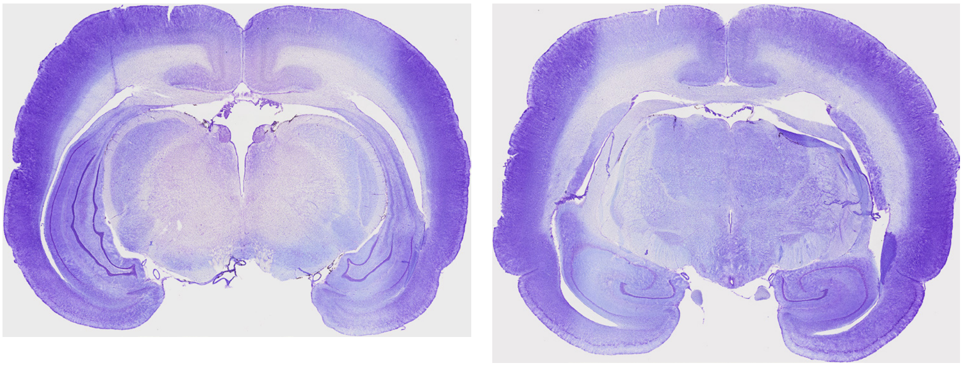

Residual HPC:
Left: 59.11 $mm^3$
Right: 32.32 $mm^3$

TS0101

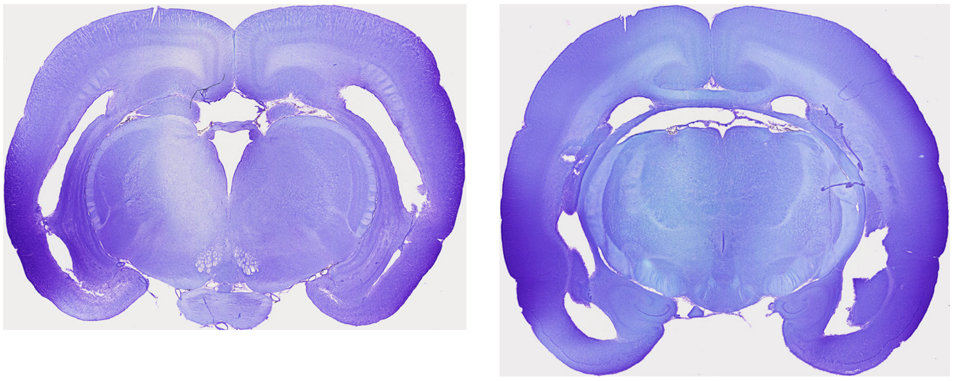

Residual HPC:
Left: 6.08 $mm^3$
Right: 5.74 $mm^3$

TS0102

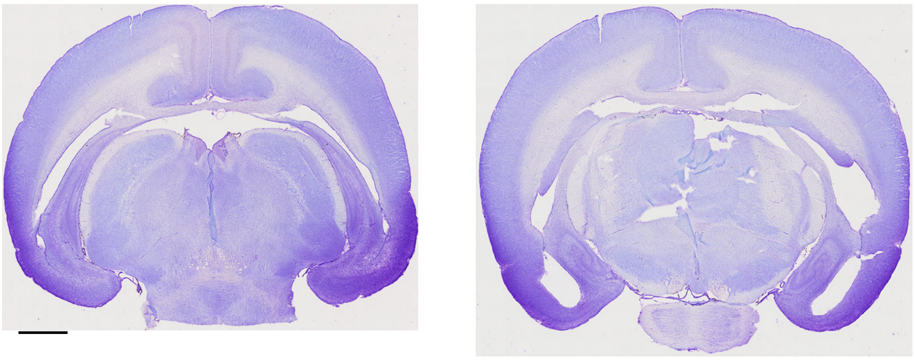

Residual HPC:
Left: 4.07 $mm^3$
Right: 3.61 $mm^3$

Figure 5-3

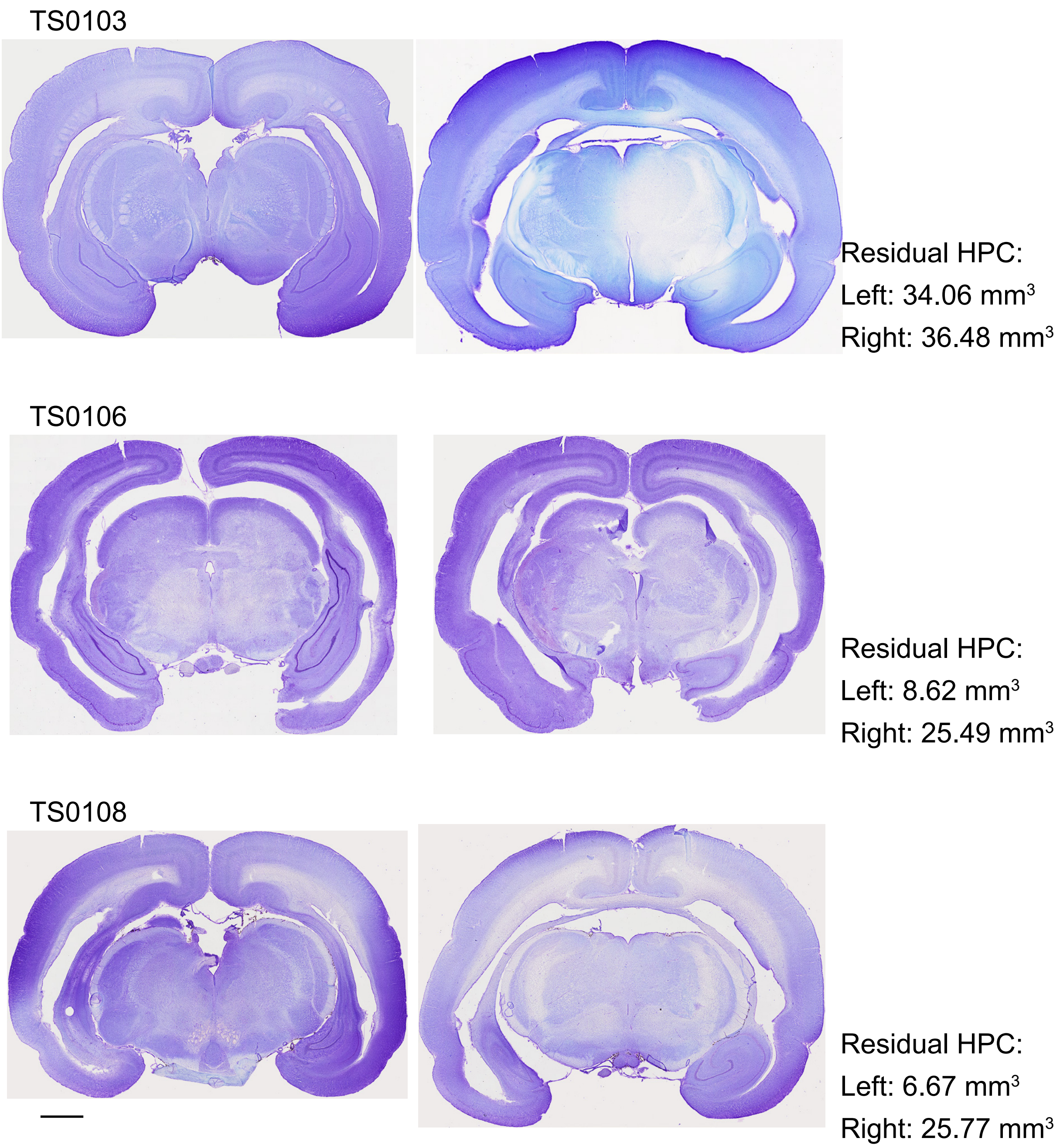


**Extended figure 5**. Representative histology showing hippocampal lesions in each tree shrew. Brain sections of similar levels from each animal are displayed. Volumes of residual hippocampal tissue in each hemisphere are indicated on the right. Scale bar: 2 mm.
